# Expression of *E. coli* glycogen branching enzyme in an *Arabidopsis* mutant devoid of endogenous starch branching enzymes induces the synthesis of starch-like polyglucans

**DOI:** 10.1101/019976

**Authors:** Laura Boyer, Xavier Roussel, Adeline Courseaux, Ofilia Mvundza Ndjindji, Christine Lancelon-Pin, Jean-Luc Putaux, Ian Tetlow, Michael Emes, Bruno Pontoire, Christophe D’Hulst, Fabrice Wattebled

## Abstract

Starch synthesis requires several enzymatic activities including branching enzymes (BEs) responsible for the formation of *α*(1→6) linkages. Distribution and number of these linkages are further controlled by debranching enzymes (DBEs) that cleave some of them, rendering the polyglucan water-insoluble and semi-crystalline. Although the activity of BEs and DBEs is mandatory to sustain normal starch synthesis, the relative importance of each in the establishment of the plant storage polyglucan (i.e. water-insolubility, crystallinity, presence of amylose) is still debated. Here, we have substituted the activity of BEs in *Arabidopsis* with that of the *Escherichia coli* glycogen branching enzyme (GlgB). The latter is the BE counterpart in the metabolism of glycogen, a highly branched water-soluble and amorphous storage polyglucan. GlgB was expressed in the *be2 be3* double mutant of *Arabidopsis* that is devoid of BE activity and consequently free of starch. The synthesis of a water-insoluble, partly crystalline, amylose-containing starch-like polyglucan was restored in GlgB-expressing plants, suggesting that BEs origin only have a limited impact on establishing essential characteristics of starch. Moreover, the balance between branching and debranching is crucial for the synthesis of starch, as an excess of branching activity results in the formation of highly branched, water-soluble, poorly crystalline polyglucan.

## INTRODUCTION

Amylopectin is one of the two homopolymers of glucose that constitute native starch in plants and algae. Like glycogen found in bacteria, fungi and animals it is composed of glucosyl residues linked by α(1→4) O-glycosidic bonds and branched through α(1→6) linkages. However, the degree of branching of amylopectin does not exceed 5-6%, while it usually reaches 8 to 10% in glycogen (Buleon *et al*., 1998, Roach *et al*., 2012). Moreover, the distribution of α(1→6) linkages is different in both polyglucans. On the one hand, in glycogen, branch points are regularly distributed in the macromolecule, leading to the formation of a spherical homogenous structure with limited size (Melendez-Hevia *et al*., 1993). On the other hand, amylopectin exhibits a heterogeneous molecular organization in which short linear segments are clustered and branch points form amorphous lamellae. Since the initial proposition of cluster organization of amylopectin (French, 1972, Robin *et al*., 1974), several models of cluster pattern have been proposed (Bertoft, 2013). Hizukuri’s model suggests that clusters follow one another linearly. The clusters are linked together by longer glucans that expand from the preceding structure to the following (Hizukuri, 1986). More recently, Bertoft has proposed an alternative model in which clusters are linked side by side to a long, linear, glucan backbone (Bertoft, 2004). In Bertoft’s model, the formation of a new cluster does not depend on the synthesis of a previous one. Regardless of which amylopectin model is considered, these differences between amylopectin and glycogen lead to different properties for both polymers. Amylopectin is water-insoluble and partly crystalline whereas glycogen is water-soluble and amorphous.

The biosynthesis of these polysaccharides involves the same set of enzymatic activities:

i. Polymerizing enzymes (i.e. starch- or glycogen-synthases) that transfer the glucose moiety of a nucleotide-sugar (being ADP-glucose or UDP-glucose) to the non-reducing end of an α-glucan by creating α(1→4) linkages (Fujita *et al*., 2012).
ii. Branching enzymes (1,4-α-glucan:1,4-α-glucan 6-glucosyltransferase; EC 2.4.1.18) that introduce the α(1→6) branch points by rearranging linear glucans. These enzymes cleave an α(1→4) linkage and then transfer the released glucan in α(1→6) position by an intermolecular or intramolecular mechanism (Tetlow, 2012).

However, about 20 years ago, it was suggested that isoamylase-type debranching enzymes (DBEs) were required for the building of the final structure of amylopectin in plants, determining insolubility and partial crystallization (Ball *et al*., 1996, Mouille *et al*., 1996, Myers *et al*., 2000). This idea was further supported and confirmed by several studies conducted with different plant species (James *et al*., 1995, Nakamura, 1996, Rahman *et al*., 1998, Delatte *et al*., 2005, Wattebled *et al*., 2005). In all cases, the lack of isoamylase-type DBEs (except the ISA3 isoform dedicated to starch degradation) caused a strong reduction of starch accumulation, and a modification of the ultrastructure of the residual starch accompanied by a decrease of the crystallinity index. Moreover, these isoamylase mutants also accumulate phytoglycogen, a water-soluble and amorphous glucan structure, which is comparable to that of the glycogen found in animals, fungi and bacteria. Compared to wild-type starch, phytoglycogen displays significant modifications of chain length distribution characterized by enrichment in very short chains (DP 3-5). The crystallization of starch clearly depends on the distribution of α(1→6) linkages, but also on chain length distribution (Pfister *et al*., 2014). Starch synthases (by elongating glucans) and isoamylases (by removing some branch points) are key enzymes that control both parameters that are crucial to define the final structure and properties of starch. Modification of plant branching enzyme (BE) activity level can affect the physicochemical characteristics of starch as well. For instance, the reduction of BE activity in potato by the antisense inhibition of both SBE A and SBE B isoforms induces the synthesis of very-high-amylose starch in potato tubers (Schwall *et al*., 2000). In rice and maize, modification of BE activity (by mutation or over-expression) alters the structure and properties of the endosperm starch (Satoh *et al*., 2003, Tanaka *et al*., 2004, Yao *et al*., 2004). Although it is obvious that BEs are mandatory for the synthesis of starch since these enzymes are the only one that create the α(1→6) linkages, their actual contribution for the determination of starch characteristics (i.e. water-insolubility, semi-crystallinity, presence of amylose, amylopectin DP_max_ at 12-13) has not been precisely evaluated so far. Recently, the synthesis of high amounts of insoluble polyglucans was reported in a quintuple mutant of Arabidopsis lacking all four DBEs plus one form of plastidial α-amylase (Streb *et al*., 2008). Thus, the necessity of DBEs *per se* for the determination of the water-insolubility and crystallinity of starch could be questioned, although the polyglucans synthesized in the above-mentioned quintuple mutant displays specific features (very high DP<10 content for instance) not observed in WT starch. Therefore, the function of BEs appears to be crucial in establishing the intrinsic structural features of starch although crystallinity of the polyglucan synthesized in the Arabidopsis quintuple mutant was not reported in (Streb *et al*., 2008). Branching enzymes are classified into two groups depending on their amino-acid sequences: group I or B-family and group II or A-family (Burton *et al*., 1995). The *in vitro* determination of catalytic parameters showed that BEI is more active on amylose than amylopectin, in contrast to the members of the BEII group (Guan *et al*., 1993, Guan *et al*., 1994). Moreover, BEI transfers longer glucans compared to BEII (Guan *et al*., 1997). GlgB that generates α(1→6) linkages during glycogen synthesis has catalytic properties obviously distinct to that of maize BEII and BEI (Guan *et al*., 1997).

In this work, our objective was to test whether the phylogenetic and family origin of BEs was crucial for the production of water-insoluble semi-crystalline polysaccharides. The expression of maize BEI or BEII or both BEI and BEII in a *glgB-* mutant of *E. coli* resulted in the synthesis of glycogen-like but not starch-like polysaccharides (Guan *et al*., 1995). Similar results were obtained after the expression of maize BEIIa and/or BEIIb in a branching enzyme mutant of yeast (Seo *et al*., 2002). Alternatively, *E. coli* glycogen branching enzyme (GlgB) was already expressed in different plants such as potato (Kortstee *et al*., 1996, Huang *et al*., 2013) and rice (Kim *et al*., 2005) leading to modification of the structure of the synthesized polyglucans (increased branching degree and altered chain length distribution of amylopectin). However in both cases, the endogenous BEs were still active, thus preventing from the determination of the specific contribution of GlgB for the synthesis of starch.

Arabidopsis contains only two BEs involved in transitory starch synthesis and both of them belong to the BEII group (Fisher *et al*., 1996). Arabidopsis *be2 be3* double mutants are unable to create branch points during polysaccharide synthesis and are therefore starchless (Dumez *et al*., 2006). We used these plants, devoid of endogenous BEs, to express the *E. coli* GlgB glycogen branching enzyme to evaluate whether enzyme metabolic origin (i.e. glycogen versus starch) can sustain the synthesis of starch-like polyglucan. More precisely, a starch-like polyglucan could be defined as a polyglucan that is water-insoluble, partly crystalline, contain both amylopectin and amylose and in which amylopectin DP_max_ is 12-13. GlgB is involved in the synthesis of glycogen and belongs to the BEI family (Sawada *et al*., 2014). The analysis of polyglucan accumulation in *Arabidopsis* plants expressing GlgB revealed that the balance between branching and debranching activities, assuming that elongating activity is not-limiting, establishes the structure of the polysaccharide produced, regardless of the origin of the branching enzyme.

## MATERIALS AND METHODS

### *Cloning of the glycogen branching enzyme (GlgB) of* Escherichia coli

The chloroplast transit peptide of *Arabidopsis thaliana* Branching Enzyme 2 (At5g03650) was identified with the use of ChloroP 1.1 (http://www.cbs.dtu.dk/services/ChloroP/). The corresponding nucleotide sequence was reconstituted by PCR using two long and partially complementary primers, TPbe2-1 and TPbe2-2 (primer sequences and method of amplification are depicted in details in Figure S1). A fragment corresponding to the first 30 nucleotides of the 5’ end of *glgB* was then added downstream of the sequence encoding the transit peptide (Figure S1). In parallel, the coding sequence of *glgB* was amplified by PCR from genomic DNA of *E. coli* TOP10 using primers glgB-For and glgB-Rev (Figure S1). The final sequence was obtained after 20 cycles of polymerase extension of both overlapping sequences. This chimeric cDNA was subsequently amplified by PCR with primers TPbe2 for-CACC and glgB rev (Figure S1) and cloned into pENTR™/D-TOPO prior to be transferred by homologous recombination into pMDC32 using the gateway^®^ technology (Life Technologies). The sequence was checked by full-sequencing before further uses. All PCR reactions were performed using a high-fidelity DNA polymerase (Kapa Hifi from KAPABIOSYSTEM®).

### Plant growth, transformation and selection

Unless otherwise indicated, Arabidopsis plants were cultivated in growth cabinet in 16-h light: 8-h dark regime at 23°C during the day and 20°C during the night under 75% relative humidity and 100 μE.m^−2^.s^−1^ irradiance.

The *be2 be3* double mutant of *A. thaliana* (WS ecotype) (Dumez *et al*., 2006) was transformed by the floral dip method using *A. tumefaciens* strain GV3101 (Clough *et al*., 1998, Jyothishwaran *et al*., 2007) according to (Facon *et al*., 2013). Transformed plants were selected for their resistance to hygromycin B. T1 seeds were sterilized according to (Harrison *et al*., 2006) and sown on petri dishes containing 1% (w/v) Murashige and Skoog medium, 1% (w/v) agar and 20 μg.mL^−1^ of hygromycin B (Sigma-Aldrich). Plates were incubated in growth cabinet for 2-3 weeks at 22°C, 80% humidity and a specific photoperiod according to (Harrison *et al*., 2006). 98 hygromycin-resistant individuals were transferred to soil and cultivated in a greenhouse under a 16-h light: 8-h dark photoperiod (20°C during the day and 18°C at night; 100-150 μΕ.m.^−2^.s^−1^ irradiance). The presence of the transgene was confirmed by PCR amplification of the *glgB* sequence (glgB-For and glgB-Rev primers) in 22 plants displaying a leaf iodine-staining phenotype (see below) different to that of the *be2 be3* mother plant. A representative panel of eleven plants with different leaf iodine-staining phenotypes was selected and T2 seeds were collected. T2 plants were selected on solid MS medium as described above and 5 homozygous lines were identified by segregation analysis of the hygromycin-resistance trait.

### Leaf iodine staining

Plants were grown in a greenhouse under a 16-h light: 8-h dark photoperiod (20°C during the day and 18°C at night; 100-150 μΕ.m^−2^.s^−1^ irradiance). Approximately 2-week-old plants were harvested at the end of the day and immediately immersed in 70% hot ethanol. Samples were incubated under shaking at 70°C and ethanol was replaced several times until complete loss of pigments, prior to be rinsed with water and stained by a KI 1% (w/v) - I_2_ 0.1% (w/v) solution.

### Extraction and quantification of polyglucans

Leaves were harvested at the end of the day, immediately frozen in liquid nitrogen and stored at - 80°C until use. Two different protocols were then applied for polyglucan extraction.

Isolation of insoluble polyglucans (for microscopy and crystallographic analysis) was adapted from (Streb *et al*., 2008). Several tens of grams of leaves were homogenized in a buffered medium (0.1 M Na-acetate (pH 4.8); 0.05% Triton X-100, 2 mM EDTA) with a blender, and then filtered on two layers of Miracloth (Calbiochem). SDS was added to the filtrate at a final concentration of 1.5% (w/v) and the insoluble polyglucans were collected by centrifugation (20 min at 20,000 g). Then, the samples were washed 4 times with water and were sequentially filtered on nylon meshes of decreasing pore size, in the order: 100, 31 and 11 μm. The filtrates were centrifuged (10 min at 16,000 g) and the polyglucan pellet was resuspended in 20% ethanol and stored at 4°C until use.

Extraction of insoluble and soluble polyglucans (for quantification and ultrastructure analysis) was performed on 1 g of leaves using the perchloric acid method described in detail in (Streb *et al*., 2008).

The polyglucan content was measured by a spectrophotometric method (Enzytec™, R-BIOPHARM®) following the manufacturer’s instructions.

### Analysis of the ultrastructure of polyglucans

The chain length distribution (CLD) of the polyglucans was determined with 200 μg of purified material debranched with a mix of 4 U isoamylase *(Pseudomonas sp.,* Megazyme) and 2U pullulanase (*Klebsiella planticola*, Megazyme) in sodium acetate buffer (55 mM final concentration, pH3.5) incubated overnight at 42°C in a final volume of 500 μL. After 10 min at 100°C, the mix was desalted on Alltech™ Extract-Clean™ Carbograph columns (Fisher Scientific). First, columns were equilibrated with 5 mL of 25% acetonitrile and washed with 5 mL of deionized sterile water. Then, samples were loaded on the columns and subsequently rinsed with 5ml of deionized water. Sample elution was performed with 2 mL of 25% acetonitrile. The samples were lyophilized and resuspended in 250 μL of deionized water. The chain length distribution of each sample was determined by HPAEC-PAD analysis (Dionex® – PA200 CarboPac column) as fully described in (Roussel *et al*., 2013).

### Starch fractionation

Starch fractionation (separation of high and low mass polymers) was performed by size exclusion chromatography on a Sepharose CL-2B matrix. 1.2 mg of starch was dispersed in 200 μL of DMSO (100%) during 15 min at 100 °C. The polymers were precipitated by the addition of 800μL of absolute ethanol and left for 2h on ice. After centrifugation for 5 min at 10,000 g, the pellet was solubilized in 500 μL of 10 mM NaOH and loaded onto the column previously equilibrated with 10 mM NaOH (0.5 cm i.d. × 65 cm) at a flow rate of 12 mL.h^−1^. Fractions of 300 μL were collected and the glucans were detected using I_2_ / KI solution (0.1% and 1% w/v respectively).

### Protein extraction

Protein extracts were obtained from approximately 1 g of fresh leaves harvested at the middle of the day and directly frozen in liquid nitrogen. Frozen leaves were finely powdered and immersed in ice-cold 100 mM Tricine/KOH (pH 7.8) 5 mM MgCl_2_, 1 mM DTT containing 0.1 % (v/v) of protease inhibitor cocktail (ProteaseArrest™, G Biosciences). Samples were centrifuged at 16,000 g at 4°C during 10 min. The supernatant was collected and the protein content was determined by Bradford assay.

### Zymogram of branching enzyme activity

Zymograms of branching enzyme activity were obtained according to the method described by (Tetlow *et al*., 2004, Tetlow *et al*., 2008). Branching enzyme activity is detected indirectly by the stimulation of phosphorylase « a ». In the presence of a high concentration of Glc-1-P and the absence of Pi, phosphorylase « a » synthesizes linear α(1→4)-linked polyglucans that can be branched in α(1→6) if a branching enzyme is in the vicinity. Branching increases the number of available non-reducing ends and thus stimulates phosphorylase « a » polymerizing activity. This leads to the synthesis of branched polyglucans that can be detected in the form of dark brown bands in the gel after iodine staining. In brief, 5% (w/v) polyacrylamide resolving gel in 375 mM of Tris-HCl pH8.8 (non-denaturing conditions) was prepared with 3 U.mL^-1^ of rabbit phosphorylase « a » (Sigma), 1 mM of maltoheptaose (Sigma) and 1.5 mM of acarbose (Sigma-Aldrich) and was maintained at 4°C. 60 μg of proteins were mixed with a loading solution in a 20:1 ratio (loading solution: 2 mg.mL^−1^ of bromophenol blue and 50% (v/v) glycerol) and loaded onto the gel. After migration (1 h 30 min at 15 V.cm^−1^, at 4°C in 25 mM Tris / 192mM Glycine buffer pH8.5), the gel was washed twice in buffer 20 mM MES and 100 mM sodium citrate at pH6.6 at room temperature. After incubation for 2 h at 30°C in 20 mM MES and 100 mM sodium citrate buffer (pH6.6) supplemented with 50 mM of glucose-1-phosphate, 2.5 mM AMP, 1 mM EDTA, and 1 mM DTT, the gel was gently rinsed three times with water and subsequently revealed with I_2_/KI solution (0.1% / 1% (w/v).

### In vitro *assay of branching enzyme activity*

The branching enzyme activity in leaf extracts was determined according to the method described in details by (Tetlow *et al*., 2008).

### Immunoblotting

Two types of immunoblotting experiments were carried out during this work. The first was performed after 100 μg of total proteins from a cell extract were separated by SDS-PAGE (7.5% acrylamide and 0.1% (w/v) SDS). After migration, proteins were transferred to a nitrocellulose membrane (Whatmann) with a transfer buffer (0.025 M Tris; 0.192 M glycine; 20% (v/v) methanol). After transfer the membrane was incubated in a blocking solution (0.01 M Tris; 0.25 M NaCl; 1.5% (w/v) BSA) during 15 min and then incubated overnight with the primary antibody diluted at 1/5000 in the blocking solution at room temperature. Then the membrane was washed three times in TTBS (0.1 M Tris; 0.25 M NaCl; 0.1% (v/v) Tween 20). The membrane was incubated for 2 h with a 1/30000 dilution of the secondary antibody (anti-rabbit IgG-alkaline phosphatase; Sigma) at room temperature. The membrane was finally rinsed three times with TTBS buffer and developed with the BCIP/NTB substrate kit following manufacturer’s instructions (Invitrogen).

In the second type of immunoblotting experiment protein extracts were treated as for BE zymogram analysis described above. However, after migration, proteins were transferred to a nitrocellulose membrane (Whatmann) following the protocol described above.

The primary antibody raised against GlgB was produced in rabbit by the inoculation of a peptide specific of the *E. coli* branching enzyme (Eurogentec). Peptide sequence: NH_2_-NLYEHSDPREGYHQDW -CONH_2_ (position 355-370 of *E. coli* GlgB protein); Protein carrier: KLH.

### Transmission and scanning electron microscopy

Strips of freshly cut leaves harvested at the end of the day were fixed with glutaraldehyde, post-fixed with osmium tetroxide and embedded in Epon resin. 70 nm-thin sections were cut with a diamond knife in a Leica UC6 microtome and post-stained with periodic acid thiosemicarbazide silver proteinate (PATAg) (Gallant *et al*., 1969). Drops of dilute suspensions of purified glucans were deposited on glow-discharged carbon-coated copper grids and the preparations were negatively stained with 2% uranyl acetate. All specimens were observed with a Philips CM200 transmission electron microscope (TEM) operating at 80 kV. Images were recorded on Kodak SO163 films. In addition, drops of briefly sonicated suspensions of purified glucans were allowed to dry on freshly cleaved mica and coated with Au/Pd. Secondary electron images of the specimens were recorded with a FEI Quanta 250 scanning electron microscope (SEM) equipped with a field emission gun and operating at 2 kV.

### Wide-angle X-ray scattering

The crystallinity index of the polyglucan samples was measured by wide-angle X-ray scattering following a method described by (Wattebled *et al*., 2008).

## RESULTS

### Selection of Arabidopsis lines expressing GlgB

*glgB* from *E. coli* was fused downstream of the nucleotide sequence encoding AtBE2 (At5g03650) chloroplast transit peptide. The resulting cDNA was then cloned into pMDC32, a binary vector designed for transgene expression under the control of the CaMV 35S promoter. *be2 be3* double mutants were transformed by the floral dip method (Clough *et al*., 1998) with an *Agrobacterium tumefaciens* strain carrying the construct. Among 6000 plants tested, 98 transformants (T1), displaying hygromycin resistance, were selected on MS agar plates. 5 independent homozygous lines were selected among their progeny (T2 plants) and further analyzed for their leaf iodine-staining phenotype (Figure 1). At the end of the daytime period, while wild-type (WT) leaves stain black due to normal amounts of starch, *be2 be3* leaves stain yellow because they lack polysaccharides. By contrast, *isa1 isa3 pu1* leaves stain orange because they accumulate high amounts of phytoglycogen (Wattebled *et al*., 2008). The phenotypes of the transformants, called “β lines”, were intermediate between those of WT and *isa1 isa3 pu1* lines, suggesting that the polysaccharide synthesis was at least partially restored. However, while β20 and β6 lines stain brown, β12 stains dark brown and β1 stains dark grey, suggesting different levels of complementation. β5 is noteworthy since iodine staining is not homogeneous in that line. A grey stain was observed at the apex of some leaves, while other leaves remained yellow. It suggests a non-homogeneous expression of the transgene in that line. For clarity, only β1, β12, and β20 lines, that are representative of each phenotype, will be further presented here. Results obtained for lines β5 and β6 are available as Supporting figures (Figure S7 and Figure S8).

**Figure 1:**
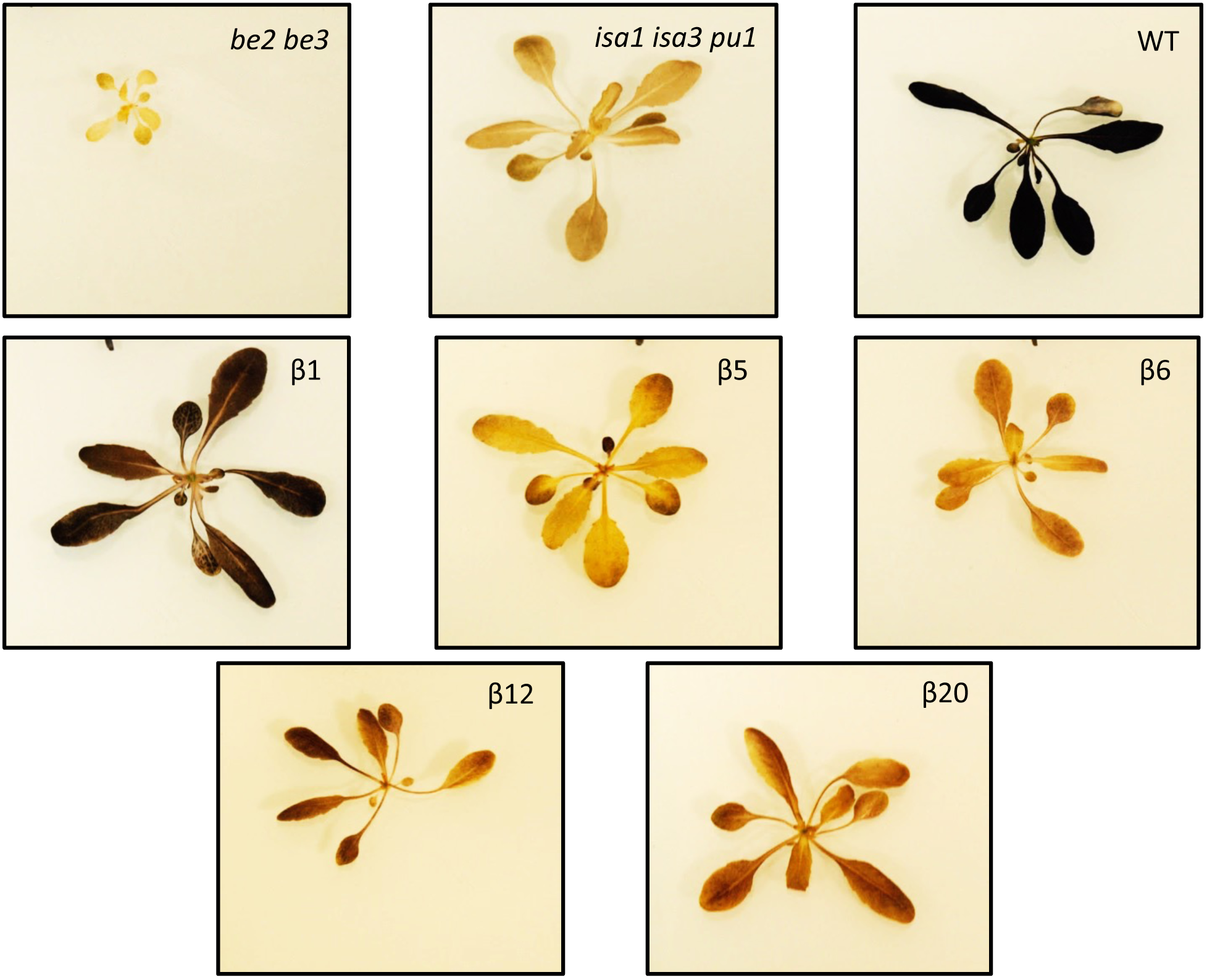
Iodine staining of individual plants harvested at the end of the day. 2-week old plants were harvested and stained with iodine solution after being destained in hot ethanol. All pictures are at the same scale. Colors and intensity of the stain is indicative of the polyglucan content and structure. Wild-type (WT) plants stain dark brown because of the synthesis of high amount of normal starch (ecotype *Wassilewkija*). The *be2 be3* plants stain yellow as a consequence of the lack of any polyglucan in this mutant (Dumez *et al*., 2006). *isa1 isa3 pu1* is a debranching enzyme triple mutant accumulating high amount of phytoglycogen (Wattebled *et al*., 2008) and therefore stains orange with iodine. β1, β5, β6, β12 and β20 are hygromycin resistant plants selected after transformation of the *be2 be3* mutant with a vector containing the *E. coli glgB* gene. Intermediate colors between those of the WT and the *isa1 isa2* mutant were obtained suggesting that polyglucan synthesis is partially restored in these transgenic lines.

The *be2 be3* double mutant displays strong growth retardation probably due to the over-accumulation of maltose (Dumez *et al*., 2006). Therefore, we analyzed the developmental phenotype of the transformants as well as *be2 be3* and wild-type plants (Figure S2). Contrary to *be2 be3* mutant plants, GlgB-expressing lines were similar to the wild type at maturity.

### Expression and activity of GlgB

The expression of GlgB in the “β” lines was confirmed by western blot using a peptide-specific anti-*E*. *coli* BE antibody raised against a 16-mer peptide of the protein (Figure 2A). As expected, while GlgB is absent in *be2 be3* and WT, a unique band of approximately 85 kDa was observed in the transgenic lines. Interestingly, each of these lines displays increasing band intensity in the order: β1, β12 and β20. Zymograms of BE activity were obtained using cell extracts prepared from leaves harvested at the middle of the light period (Figure 2B). As already described (Dumez *et al*., 2006), no BE activity is observed in the *be2 be3* mutant of Arabidopsis while WT displays one band of activity on the zymogram (Figure 2B). In contrast, a large activity band of lower mobility was found in the *E. coli* extract. BE electrophoretic profiles of the transgenic lines differ between each other as well as from those of *E. coli* and Arabidopsis. Indeed, one band of intermediate mobility (Figure 2B, a + b) is present in all extracts of the three “β” lines except in line β5 in which expression of GlgB seems extremely low and heterogenous throughout the plant (Figure 2B). Moreover, an additional band with a higher mobility than that of Arabidopsis BE is visible in line β20 (Figure 2B, c). Band intensities also vary among “β” lines, being far greater in β12 and β20 extracts compared to β1 (Figure 2B).

**Figure 2:**
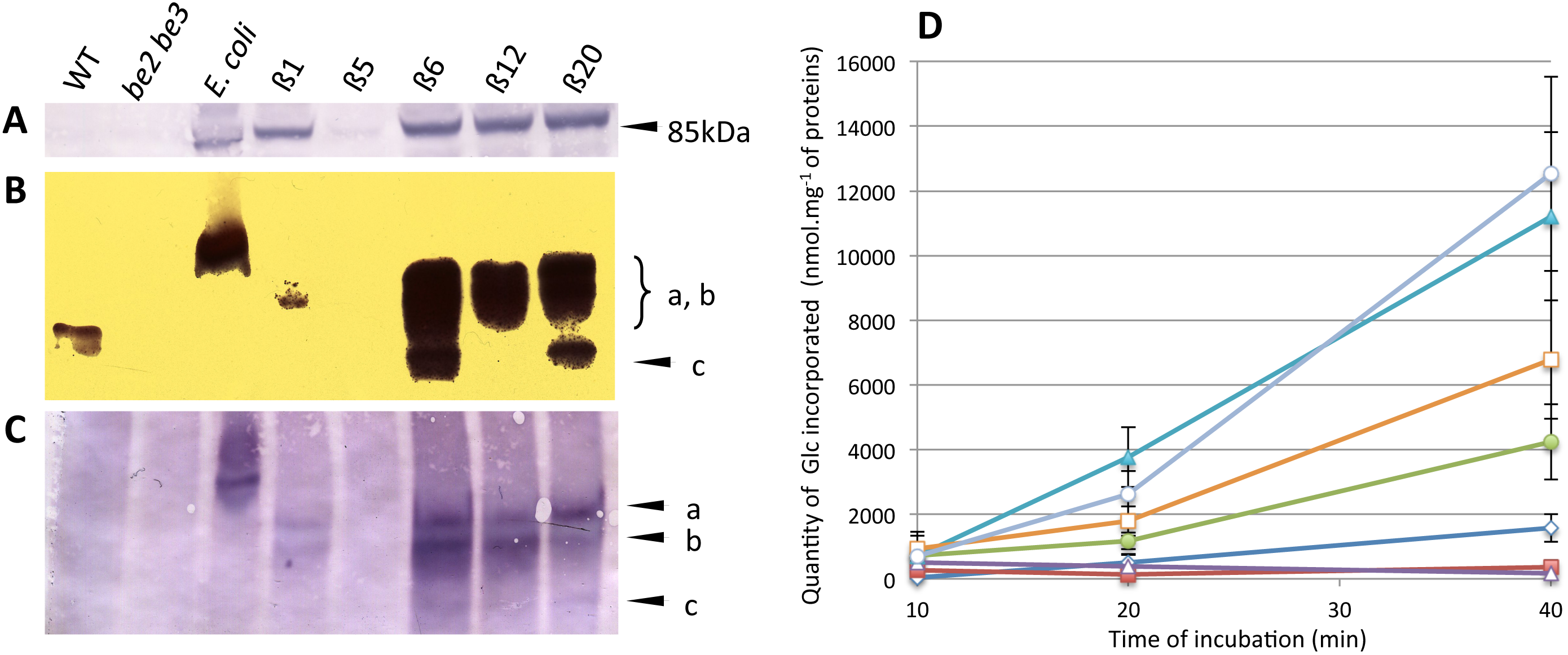
Expression of glgB in the *be2 be3* transformed lines. (A) Immunoblot analysis performed in denaturing conditions. Protein extracts were denatured before and during migration in the polyacrylamide gel. After migration, proteins were blotted onto a nitrocellulose membrane and hybridized with a peptide-directed antibody raised against GlgB. (B) zymogram of branching enzyme activity. The polyacrylamide gel contains DP7 maltooligosaccharides and phosphorylase “a”. Branching enzyme activity was revealed by incubating the gel for 2 hours in a phosphorylase “a” stimulating buffer. Bands of branched polyglucans were detected by soaking the gel in iodine solution. (C) Immunoblot analysis performed in non-denaturing conditions: protein extract preparation and migration in the gel was carried out as in (B). After migration, proteins were treated as in (A). WT: wild type of *A. thaliana* (*Wassilewskija* ecotype); β1, β12, β20: transgenic plants expressing GlgB; *be2 be3:* branching enzyme double mutant (Dumez et *al*., 2006); *E. coli:* bacterial cell extract of wild type *E. coli.* Proteins or activity bands corresponding to GlgB expressed in plants were labeled as a, b, and c. In (A), (B) and (C) all lines are from the same gel and blots. (D) *in vitro* assay of branching enzyme activity. Protein extracts were incubated for 10 to 40 min in a buffer containing DP7 maltooligosaccharides, phosphorylase “a” and [U-^14^C]Glc-1-P (7.4 kBq per assay) at 50 mM final concentration. The mean of incorporated Glc into branched polyglucans was calculated for 3 independent assays and plotted against the time of incubation. Vertical lines stand for standard deviation. Open diamonds: WT extract; closed squares: *be2 be3* mutant; closed circles: β1; open triangles: β5; closed triangles: β6; open squares: β12; open circles: β20.

Correlations between zymogram activities and *glgB* expression in the transgenic lines were sought by western blot. Leaf or cell protein extracts were separated by polyacrylamide gel electrophoresis in non-denaturing conditions similar to those used in the zymogram assay. As expected, no cross-reaction could be seen in *Arabidopsis* WT and *be2 be3* samples. One band was observed in the *E. coli* extract, the mobility of which corresponded to the activity detected on the zymogram (Figure 2B and 2C). Similarly, two bands corresponding to the activities observed on the zymogram were visible in the β1, β12, and β20 transgenic lines (Figure 2C, a and b). However, the intensity of the two bands in line β1 was lower than that of the two other transgenic lines. A third band of higher mobility was also clearly visible in line β20 (Figure 2C, c) and may correspond to the high mobility activity observed on the zymogram.

Zymogram and immunoblotting assays suggest that different levels of BE activity can be found in the lines tested in this work. Thus, BE activity was measured *in vitro* by the phosphorylase « a » stimulating assay using ^14^C-labeled Glc-1-P (Figure 2D). Leaf extracts were incubated for 10, 20 or 40 min at 30°C and Glc-1-P consumption was plotted against the incubation time (Figure 2D). This assay confirmed that BE activity increased in the “β” lines, in the order: β1, β12 and β20. An extremely low level of activity was detected in the *be2 be3* mutant of Arabidopsis (phosphorylase « a » activity is not stimulated because of the lack of BE activity in this line) while WT shows significant incorporation of Glc into polyglucan after 40 min of incubation.

### Polyglucan accumulation in the glgB expressing lines

Iodine-staining phenotypes of the transgenic lines expressing *glgB* suggest that starch and/or water-soluble polysaccharide synthesis is at least partially restored despite the lack of endogenous branching enzymes. These polyglucans were assayed after extraction from leaves harvested at the end of the 16-h light period (Figure 3). Wild-type Arabidopsis plants accumulate insoluble polysaccharides and only tiny amounts of non-methanol-precipitable sugars (i.e. glucose and maltooligosaccharides), while methanol-precipitable polysaccharides could not be detected (Figure 3). Interestingly, line β1 displayed the highest amount of insoluble polyglucans among transgenic lines while it also showed the lowest BE activity (Figure 2 and Figure 3). In line with this, insoluble polysaccharide contents were lower in both β12 and β20, in contrast to the levels of BE activity observed in these plants. Interestingly, methanol-precipitable glucans were detected at significant levels in β12 and β20 compared to β1. Non-methanol-precipitable glucans were found at a relatively high level in β20 (about 1 mg.g^−1^ of fresh leaves) compared to the other lines (0.35 mg.g^−1^ in β12; 0.2 mg.g^−1^ in β1; 0.05 mg.g^−1^ in WT). In the same culture conditions (16 h light), the *be2 be3* double mutant accumulated very high amounts of maltose (up to 20 mg.g^−1^ of fresh weight) (Dumez *et al*., 2006). Finally, note that the glucans accumulated in the different lines analyzed in this work were degraded in the night as shown by leaves iodine staining (Figure S3).

**Figure 3:**
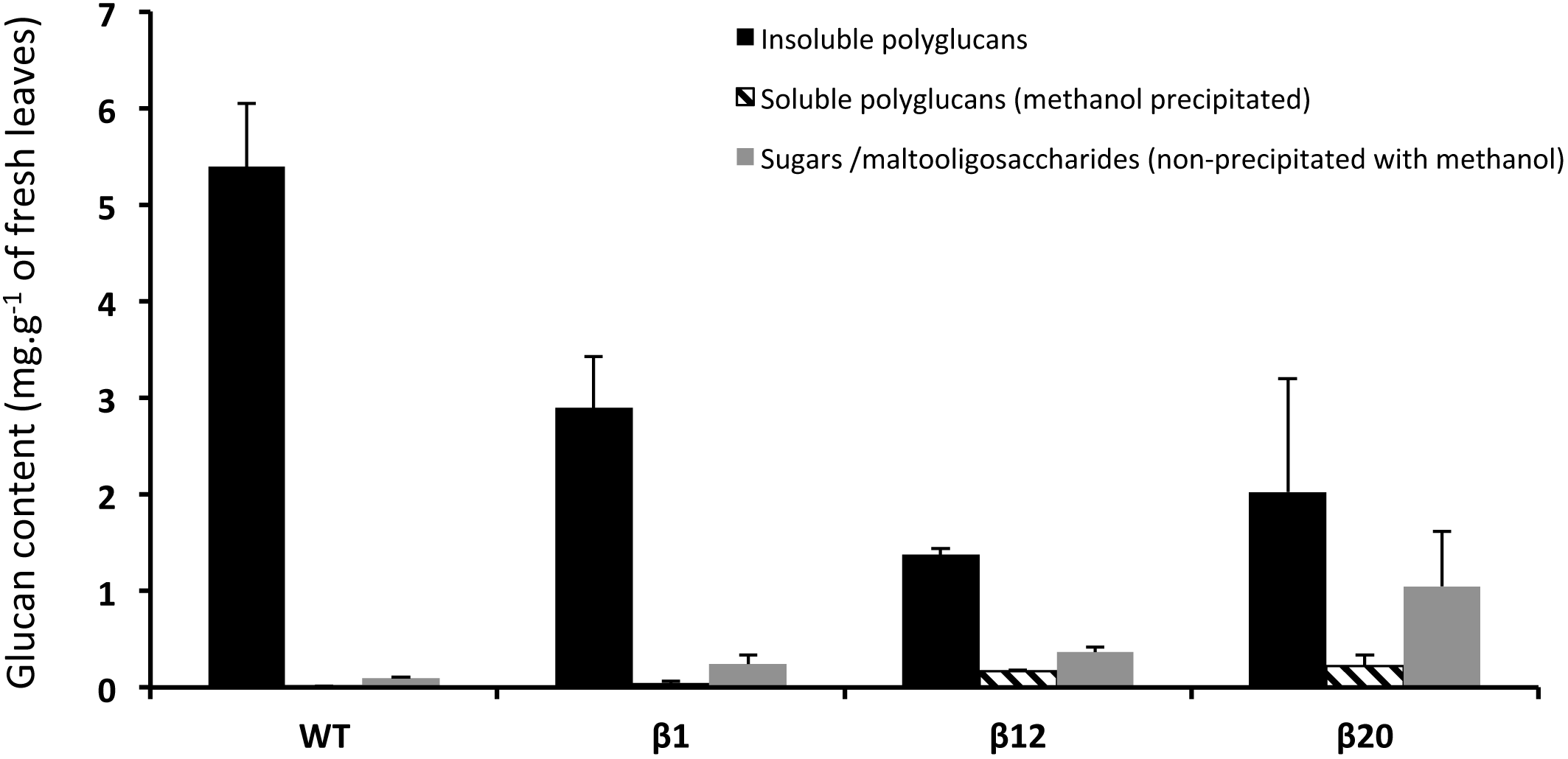
Leaf glucan contents. Glucans were extracted from leaves of 3-week old plants by the perchloric acid protocol. Leaves were harvested at the end of 16-h light period. Insoluble and soluble polyglucans and sugars/maltooligosaccharides were assayed by a spectrophotometric method. The averages of three independent cultures are presented. WT: wild type *WS* ecotype; β1, β12, β20: transgenic plants expressing the *E. coli glgB* gene. Vertical thin bars are the standard deviation of three independent biological replicates.

### Observation of polyglucans accumulation in leaf chloroplasts

The accumulation of polyglucans was directly observed *in planta* by transmission electron microscopy (TEM - Figure 4). Ultrathin sections of leaves were observed after positive staining of the polyglucans with PATAg. Flat and smooth starch granules of approximately 1-3 μm were observed in WT (Figure 4A), as already described in the literature. By contrast, no starch granules were detected in the *be2 be3* double mutant, in agreement with the phenotype already described for this line (Figure 4B) (Dumez *et al*., 2006). The *isa1 isa3 pu1* triple mutant was included in this analysis as a starchless phytoglycogen-accumulating line. Phytoglycogen was visible in the form of very small and polydisperse particles homogeneously distributed in the stroma of the chloroplasts (Figure 4C). Different types of particles were observed in the three lines expressing *glgB*. Lines β1 and β12 accumulated 0.2 - 3 μm-large and irregularly shaped particles (Figure 4D and 4E, respectively), while β20 contained significantly smaller and highly polydisperse particles (Figure 4F). The larger objects could reach 1 μm in size but most particles were smaller than 0.2 μm. The image of β20 is thus similar to that of *isa1 isa3 pu1* (Figure 4C) although no large aggregates were observed in the images of the triple DBE mutant. The surface of the particles was irregular and rather rough in β1 while it was smoother in β12. In both specimens, the fact that the larger particles may correspond to the aggregation of smaller units cannot be excluded.

**Figure 4:**
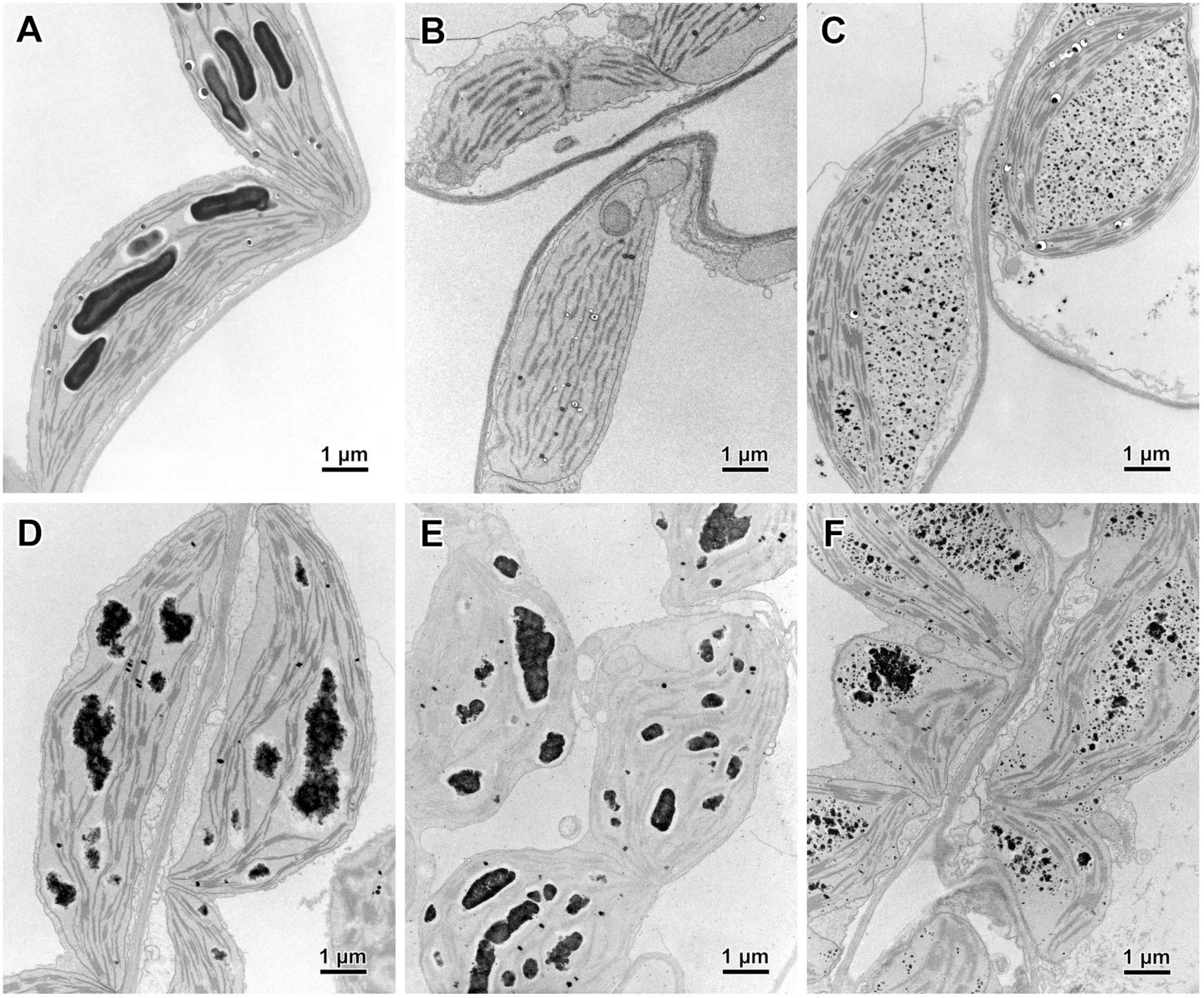
Transmission electron microscopy of leaf chloroplasts. Leaves from plants were harvested at the end of the day, cut in small strips and immediately fixed with glutaraldehyde in a cacodylate buffer and stained with PATAg to reveal glucans. A: wild type from *Wassilewskija* ecotype of *A. thaliana;* B: *be2 be3:* branching enzyme double mutant; C: *isa1 isa3 pu1:* debranching enzyme triple mutant accumulating phytoglycogen (Wattebled *et al*., 2008); D: β1, E: β12, F: β20: *be2 be3* transgenic plants expressing the bacterial *glgB* gene.

### Morphology of purified insoluble and soluble polyglucans

The insoluble polyglucans were observed by scanning electron microscopy (SEM) after extraction from leaves and purification (Figure 5). In WT, starch granules appeared as polygonal flat particles with a smooth surface, as usually observed for Arabidopsis starch granules (Figure 5A). Their size typically ranged between 2 and 5 μm. The identification of individual particles in extracts from the transgenic lines was rather difficult as the images showed that the particles were highly polydisperse and with very irregular shapes (Figure 5B-5D). Artefactual particle aggregation could not be excluded since, during purification, the fraction were extensively washed using repeated centrifugation. Local concentration of the smallest particles upon drying on the supporting mica was also unavoidable as they were more easily subject to the capillary effects of evaporating water. Nevertheless, the images were in fairly good agreement with what was expected from the TEM observation of ultrathin sections of leaves (Figure 4D-4F), with the particle average size clearly decreasing with increasing GlgB activity. In particular, the particles in β1 exhibited tortuous shapes and a rather rough surface (Figure 5B), while two polyglucan populations were observed in β20 (Figure 5D): one with an average size ranging from 500 nm to 1 μm, quite similar to those observed in β12 (Figure 5C), and one containing smaller particles (< 200 nm) whose morphology was hardly resolved by SEM (inset Figure 5D).

**Figure 5:**
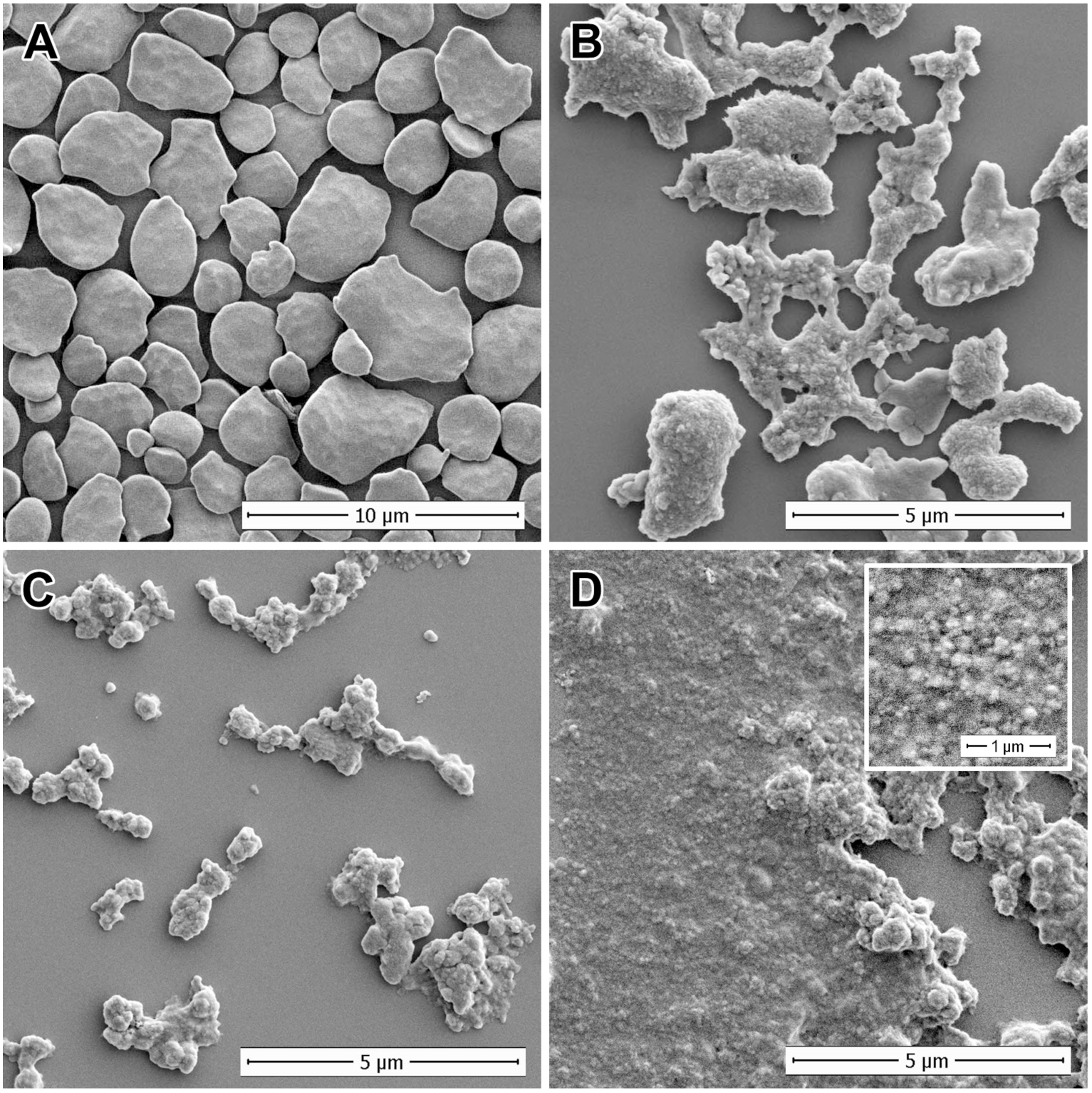
Scanning electron microscopy of purified insoluble polyglucans. Insoluble polyglucans were extracted and purified from leaves of 3-week-old plants harvested at the end of the day of 16-h light / 8 h dark cycles. A: wild-type *Wassilewskija* ecotype; B: β1; C: β12; D: β20 (inset: magnified view of the smallest particles).

Higher resolution images of the insoluble polyglucans were recorded by TEM of negatively stained preparations. As the particles extracted from β1 were too large to be observed with this technique, only particles extracted from β12 and β20 are shown in Figure 6. Moreover, only the smallest of these insoluble particles could be properly stained and observed with sufficient detail. Consequently, the images illustrate the morphology of this fraction of particles but may not represent that of the larger objects.

**Figure 6:**
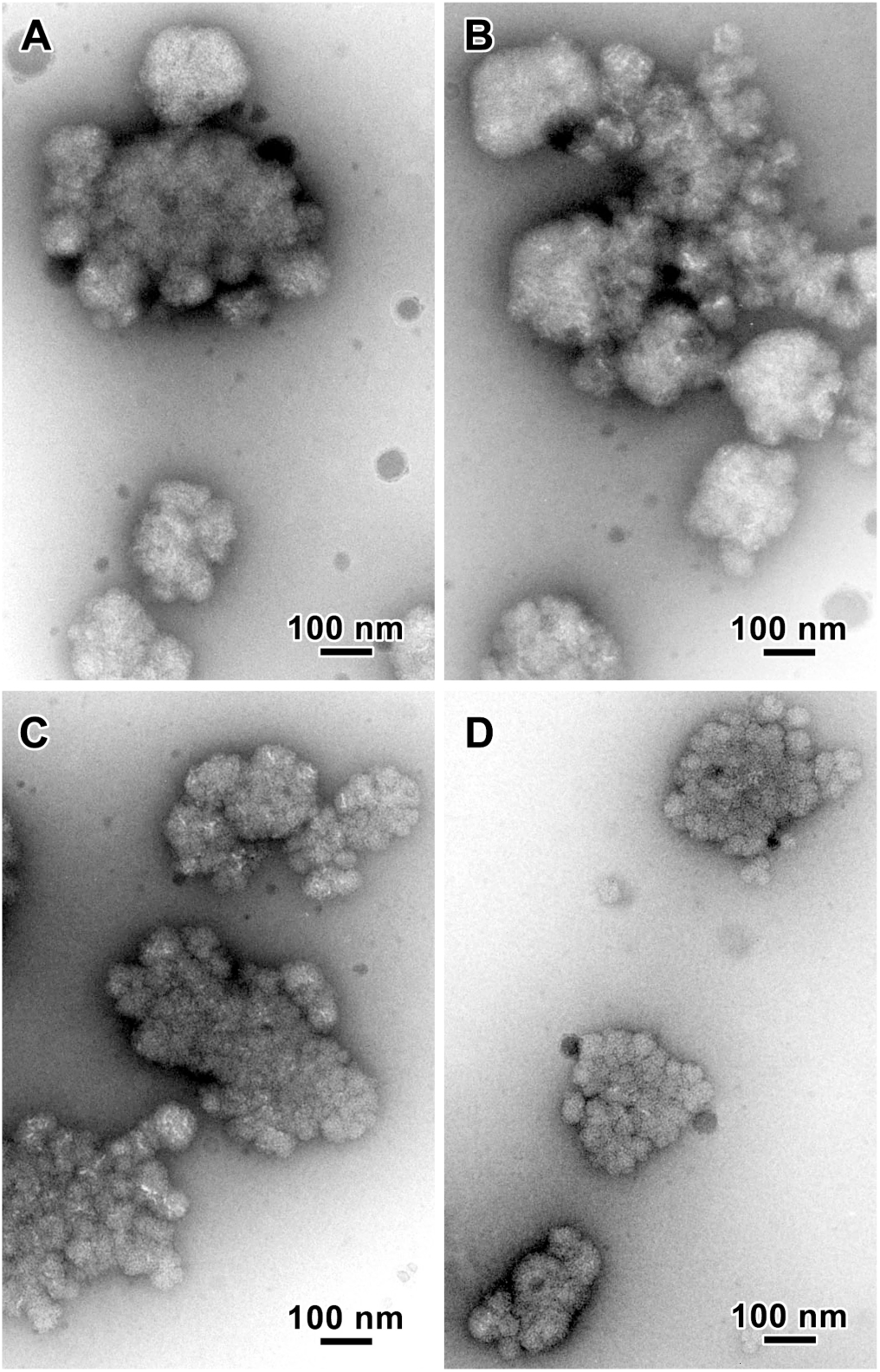
Morphology of particles purified from the insoluble polyglucans of the GlgB-expressing lines. TEM images of negatively stained particles from the insoluble fractions of β12 (A, B) and β20 (C, D).

The insoluble polyglucans in β12 often appeared as aggregates of smaller particles with various shapes and sizes (Figure 6A and 6B). 200 nm bulky and rather smooth spheroidal particles could be recognized, often in contact with multilobular less well-defined material. The particles in β20 are more flat and exhibit a clear multilobular aspect (Figure 6C and 6D). Their average size is still around a few hundreds nanometers but they seem to be composed of 50-70 nm subunits. Although it is difficult to see if the larger particles are aggregates of individual subunits formed upon centrifugation and/or drying, the morphology resembles that of the so-called α-particles observed in liver glycogen and constituted of smaller β-subunits (Ryu *et al*., 2009, Sullivan *et al*., 2012).

TEM images of the polyglucans isolated from the soluble fractions from β12 and β20 are shown in Figure 7. In both cases, 20-30 nm particles are observed (Figure 7A and 7B) whose aspect and size distribution are very similar to those of maize phytoglycogen (Figure 7C).

**Figure 7:**
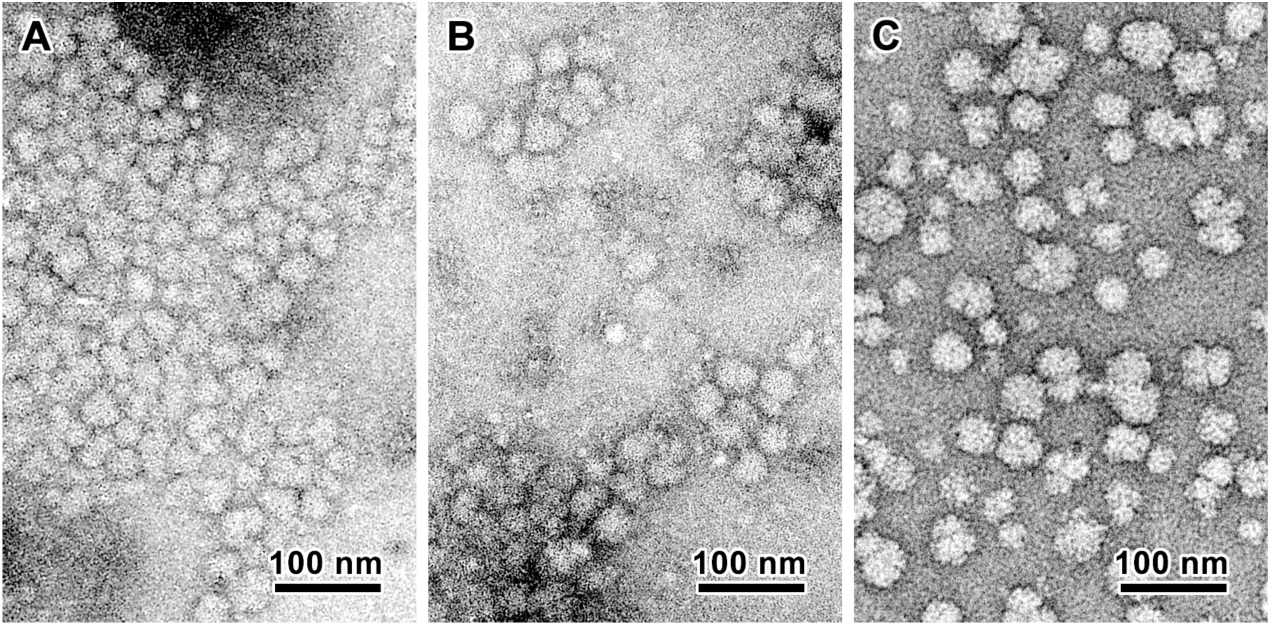
Morphology of particles purified from the soluble polyglucans isolated from the GlgB-expressing lines. TEM images of negatively stained particles from the water-soluble polyglucans isolated from β12 (A) and β20 (B). The image of maize phytoglycogen (C) is given for comparison.

### Ultrastructure of insoluble and soluble polyglucans

The ultrastructure of insoluble and soluble polyglucans was determined and compared to that of reference samples. First, insoluble polyglucans were fractionated by size exclusion chromatography on Sepharose CL-2B® (Figure 8). WT starch shows a regular elution profile exhibiting two peaks (Figure 8A). The first peak corresponds to high-mass amylopectin (between 10 and 13 mL) while the second broader peak corresponds to the low-mass amylose (between 16 and 27 mL). The elution profile of insoluble polyglucans for line β1 also contains two distinct peaks corresponding to high-mass (same elution volume as WT amylopectin) and low-mass material (elution volume equivalent to that of WT amylose), respectively (Figure 8B). However, the peak of low-mass material is much higher and slightly shifted towards lower masses (to the right on the diagram) compared to WT amylose. Moreover, the *λ*_max_ of the iodine-polysaccharide complex for high-mass material in line β1 is 10 nm higher compared to that of WT amylopectin. This suggests that this material is composed, on average, of longer-chain glucans than WT amylopectin. β12 and β20 insoluble polyglucans display more or less the same elution profile (Figure 8C and 8D, respectively). A unique large peak of high-mass material is observed, the elution profile of which is equivalent to that of WT amylopectin. Contrary to WT and β1 insoluble polyglucans, no distinct peak of low-mass material was seen in β12 and β20 lines. In addition, the *λ*_max_ of the high-mass material in these two lines was decreased by 10 nm compared to WT amylopectin, suggesting that the length of the glucans is shorter on average (Table 1).

**Figure 8:**
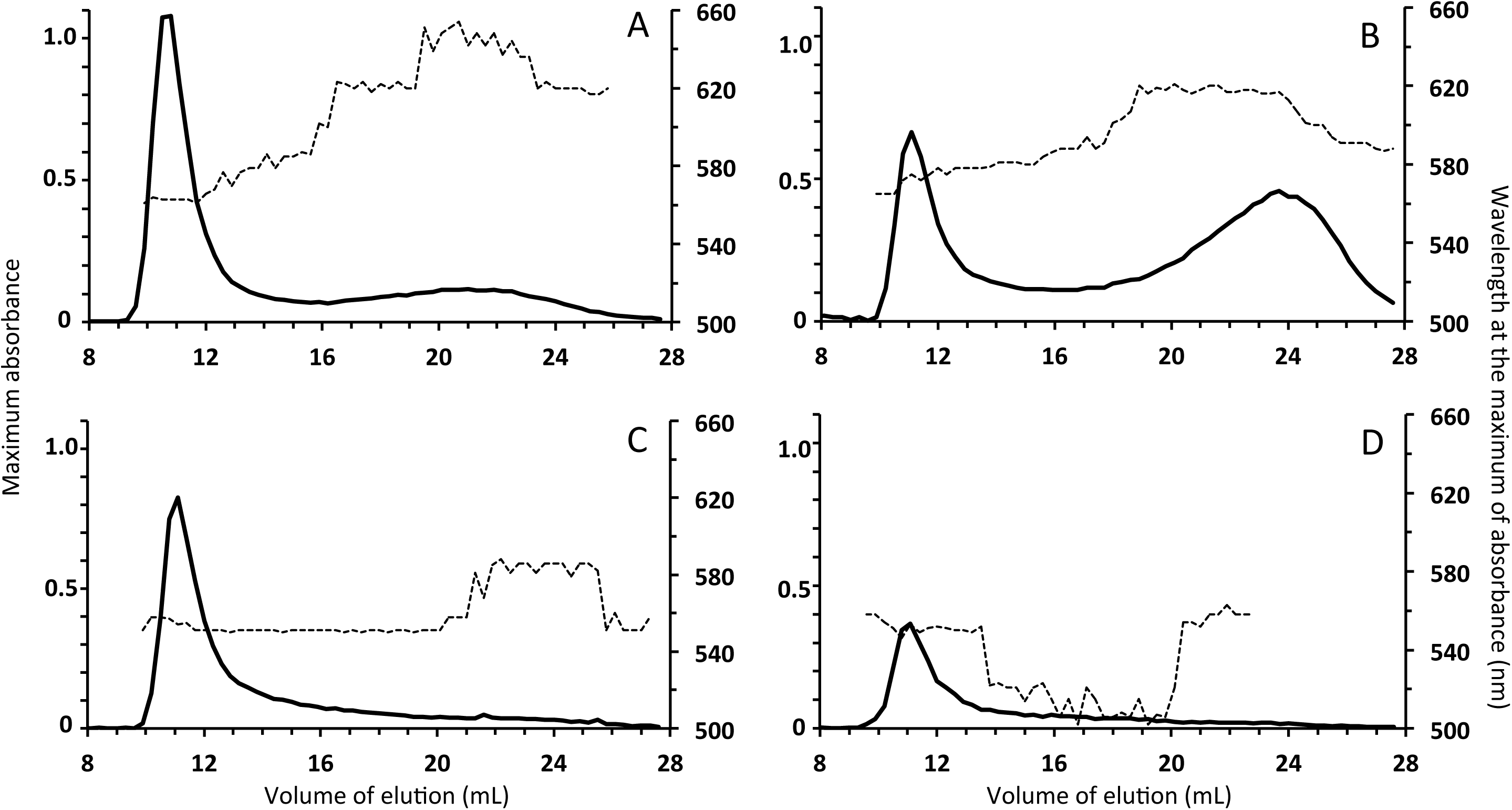
Fractionation of insoluble polyglucans by size exclusion chromatography. Purified insoluble polyglucans were dispersed in DMSO, solubilized in NaOH 10 mM and loaded onto a Sepharose® CL-2B matrix. Elution was conducted in NaOH 10 mM at a rate of 12 mL.h^−1^. 300 μL fractions were collected and subsequently analyzed by iodine spectrophotometry. Thick continuous lines are the maximum absorbance of the iodine-polyglucan complexes (left Y-axis). Dashed lines indicate the wavelength (nm) of the iodine-polyglucan complex at the maximum of absorbance (right Y-axis). X-axis is the volume of elution in mL. A: WT; B: β1; C: β12; D: β20.

**Table 1:** The branching degree was calculated according to (Szydlowski *et al*., 2011). The λ_max_ of the iodine-amylopectin complex was determined after insoluble polyglucans fractionation by SEC on CL-2B matrix (). It was determined at the fraction of maximum absorbance. The crystallinity index and allomorph were determined by wide-angle X-ray scattering. All samples were of B-type. The values are the means of two independent assays. In the case of the *isa1 isa3 pu1* triple mutant, the branching degree was determined for phytoglycogen. NA = not available.

| Sample | WT | $\beta 1$ | $\beta 12$ | $\beta 20$ | *isa1 isa3 pul* |
| --- | --- | --- | --- | --- | --- |
| Branching degree (%) | 5.1 | 6.1 | 6.8 | 7.3 | 8.1 |
| $\lambda_{\max}$ of amylopectin (nm) | 563 | 572 | 553 | 552 | NA |
| Crystallinity index (%) | 33 | 27 | 20 | 13 | NA |

The crystallinity index of the insoluble polyglucans was determined by wide-angle X-ray scattering. The analyses were performed on two samples of polyglucans extracted from two plant preparations cultivated independently (Table 1). All samples are of B-type allomorph. The WT starch granules have a crystallinity index of 33%, in agreement with previous reports on Arabidopsis starch (Wattebled *et al*., 2008). The β1 and β12 insoluble polyglucans display a lower crystallinity index (27 and 20%, respectively), while the β20 insoluble glucans have the lowest crystallinity index (13%) among those analyzed in this work.

The chain length distribution (CLD) profile of the insoluble polyglucans was established after debranching with a mixture of bacterial isoamylase and pullulanase. The resulting linear glucans were separated and quantified by HPAEC-PAD (Figure 9). The profile of β1 polyglucans has a DP max of 12-13 glucosyl residues identical to WT starch (Figure 9A and 9C). However, the fraction of DP 5-18 glucans is higher and the fraction of DP 21-40 is lower in β1 compared to WT. The higher fraction of short glucans (DP<13) that do not interact with iodine may explain the increase of the *λ*_max_ of the iodine-polyglucan complex observed in this sample (Table 1). The β12 insoluble polyglucan has a profile similar to that of β1 although the fraction of DP 5-11 glucans is higher, especially at DP 6-7 (Figure 9A). In β20, the DP_max_ is at DP 7 and the profile strongly resembles that of the insoluble glucan synthesized by the *Arabidopsis isa1 isa3 pu1* triple mutant (Figure 9B) apart from for a strong depletion of DP 3-4 glucans in β20 that is probably due to the presence of DBE activity in that line compared to the triple DBE mutant.

**Figure 9:**
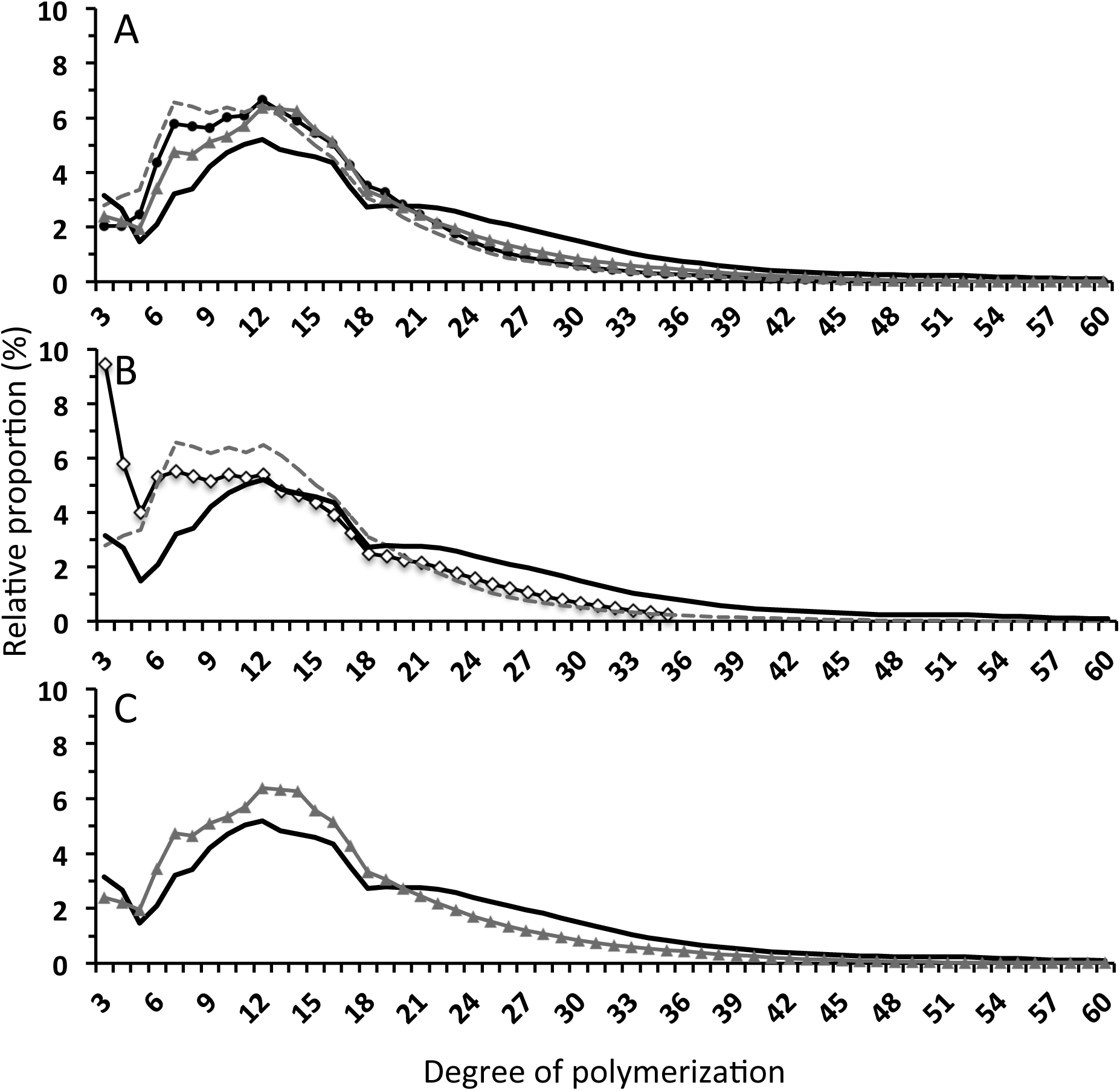
Chain length distribution of insoluble polyglucans. Purified insoluble polyglucans were digested with a mix of bacterial debranching enzymes. The corresponding linear glucans were separated by high-performance anion exchange chromatography and detected by pulsed amperometric detection (HPAEC-PAD). The fraction of each DP (from 3 to 60 glucose residues) is expressed in % of the total DP presented on the profile. (A) WT: continuous black line; β1: grey triangles; β12: closed circles; β20: discontinuous grey line. (B) Comparison of the profiles of the WT (continuous black line) and β20 (discontinuous grey line). The profile of the *isal isa3 pul* triple mutant already published in (Wattebled *et al*., 2008) is given (open diamonds). (C) Comparison of the profiles of the WT (continuous black line) and β1 (grey triangles).

The branching degree of the insoluble polyglucans was estimated according to the method described in (Szydlowski *et al*., 2011) (Table 1). WT, β1, β12 and β20 polyglucans had branching degrees of 5.1, 6.1, 6.8 and 7.3%, respectively. The branching degree of the phytoglycogen of the *Arabidopsis isa1 isa3 pu1* triple mutant was estimated at 8.1%, a value very close to that previously reported (Szydlowski *et al*., 2011).

The CLD profiles were also established for the soluble polyglucans isolated from the transgenic lines. The corresponding profiles were compared to that of rabbit liver glycogen (Figure S4). In all cases, DP 7 chains were the most abundant among those analyzed. Although the profiles of the transgenic lines were all similar, they were different to that of glycogen with a significant excess of DP 5-13 chains and a deficit in DP > 17 chains.

## DISCUSSION

The suppression of both BE2 and BE3 in *Arabidopsis* resulted in the complete loss of starch branching enzyme activity (Figure 2B and 2D) and the loss of starch synthesis and was accompanied by the accumulation of maltose and plant growth retardation (Dumez *et al*., 2006). Other starch metabolizing enzymes were not affected by the mutations. Thus, the *be2 be3* double mutant represents an excellent model to investigate the ability of GlgB to restore starch or starch-like polyglucan synthesis in Arabidopsis.

The expression of *glgB* in *A. thaliana* was successfully achieved after transformation of the corresponding cDNA into *be2 be3* plants. The corresponding protein was detected by immunoblot analysis of leaf extracts of several independently transformed plants (Figures 2A and 2C) and the branching activity of the enzyme was also detected (Figures 2B and 2D). Different levels of expression were obtained, although all plants analyzed in this study were homozygous for the transgene. Indeed, the level of BE activity was much higher in β20 compared to β12, and most notably compared to β1 which had the lowest BE activity as measured by zymogram and *in vitro* radioactive assay (Figures 2B and 2D). These levels of activity were correlated with GlgB protein contents. Immunoblot analysis performed under denaturing or non-denaturing conditions (Figures 2A and 2C respectively) showed that the content of GlgB is higher in β20 than in β12 and even more than in β1, which is in good agreement with the level of activity assayed in each line. Interestingly, immunoblot analysis performed under non-denaturing conditions (Figure 2C) revealed several bands reacting with the anti-GlgB antibody in the transgenic lines. Two major bands (Figure 2B bands a and b) were detected in all three lines and a third band (Figure 2B band c) of higher mobility but lower intensity was specifically observed in β20. Two additional lines were analyzed during this work: β5 and β6. Corresponding results are presented as Supporting data in the form of Figures S6, S7 and S8 and confirm the results presented above. Line β5, which has a very low GlgB activity, displays a phenotype similar to that of β1. Conversely line β6, that has high GlgB activity, is similar to β20. This result suggests that GlgB is either post-translationally modified when expressed in *Arabidopsis* or is involved in the formation of protein complexes (homo or heteromeric complexes) with different electrophoretic mobility (Tetlow *et al*., 2004, Makhmoudova *et al*., 2014).

Although GlgB was expressed and active in all transgenic lines, the phenotype of polyglucan accumulation was different from one line to another. Indeed, albeit displaying the lowest GlgB expression level, β1 accumulated polyglucans with a structure close to that of WT starch. These polyglucans had a granular morphology and individual particles were observed in the stroma of the chloroplast (Figure 4). These granules had a size similar to that of WT starch although their surface appeared to be rougher. Smaller PATAg-stained particles, with a size well below 1 μm, were also dispersed throughout the stroma of β1 chloroplasts. Moreover, the β1 insoluble polyglucan fraction was partially crystalline (27% compared to 33% for WT starch) and was composed of both high-mass amylopectin-like material and low-mass amylose-like material (Figure 8). Further, the CLD profile after debranching was similar to that of WT starch although it was enriched in DP 5-19 chains (with a DP_max_ at 12 glucose residues such as in WT starch) and depleted in DP 21-40 chains (Figure 9). As a whole, the β1 polyglucan can be considered as starch-like implying that a bacterial glycogen BE can sustain the synthesis of such specifically ordered polysaccharide when expressed in plants. It follows that the final distribution of the branch points within starch and consequently the properties of the synthesized polyglucan (crystallinity, water-insolubility, granular morphology, amylopectin DP_max_ at 12-13, and the presence of amylose) is not solely under the control of starch branching enzymes (although branching enzyme activity is mandatory to create α(1→6) linkages). Otherwise, expression of a bacterial glycogen-branching enzyme in the plant would have resulted in the synthesis of soluble and amorphous glycogen-like polymers. Nevertheless, the polyglucan that accumulates in β1 can not be regarded as true WT starch and several hypotheses can be proposed which could explain why this is the case. Firstly, GlgB is expressed in *Arabidopsis* under the control of the constitutive 35S promoter. Consequently, GlgB is likely expressed linearly during the day and the night, which may influence the final structure of the synthesized polyglucan. However, the expression profiles of the endogenous BEs (BE2 and BE3) are not similar during the day/night cycle (Smith *et al*., 2004) although their functions are largely overlapping (Dumez *et al*., 2006). Indeed, BE2 exhibits only slightly higher expression in the day compared to the night. BE3, expression is significantly higher during the day compared to night and displays an expression pattern that resembles that of enzymes involved in starch degradation (Smith *et al*., 2004), although, more recently, BE3 protein abundance was reported as unmodified throughout the day/night cycle (Skeffington *et al*., 2014). The second hypothesis relies to the intrinsic properties of GlgB activity. Indeed, starch BE isoforms have different catalytic properties (Tetlow, 2012). In *A. thaliana,* in contrast to other plants and to other dicots, only class-II branching enzymes (BE2 and BE3) have been described so far. However, because of its catalytic properties, GlgB can be classified in the class-I plant branching enzymes and preferentially transfers DP 6-15 chains (with a maximum for DP 10-12 chains) (Guan *et al*., 1997, Sawada *et al*., 2014). Therefore it is not surprising that the expression of a class-I BE instead of class-II BEs results in modifications in the structure of the synthesized polysaccharide compared to the wild type. Another explanation relates to enzyme complex formation. It is now well established that some of the starch metabolizing enzymes interact to form hetero-multimeric complexes (Hennen-Bierwagen *et al*., 2008, Tetlow *et al*., 2008, Hennen-Bierwagen *et al*., 2009). Branching enzymes are important components of these complexes as shown in the endosperm of cereals (Liu *et al*., 2009, Liu *et al*., 2012a, Liu *et al*., 2012b). Although it has not been shown yet, it is likely that starch-metabolizing enzymes also organize in the form of multisubunit complexes in Arabidopsis. It is highly probable that BE2/BE3 and GlgB are engaged in different protein complexes or simply that GlgB is unable to interact with other endogenous plant enzymes. This could result in the modification of other starch-metabolizing activities and consequently polyglucan structure. Even if enzymes of starch synthesis do not physically interact in Arabidopsis chloroplasts, their activities must still be interdependent in order to produce the final structure of amylopectin. Removing one or several enzymes of the pathway could alter the activity of others whose normal activity depends on the presence of the complete set of enzymes (Szydlowski *et al*., 2011, Nakamura *et al*., 2012, Abe *et al*., 2014, Brust *et al*., 2014, Nakamura *et al*., 2014). GlgB may only partially counterbalance the lack of both BE2 and BE3 in the transgenic lines, which, in turn, may alter other starch-metabolizing activities. Lastly, some starch-metabolizing enzymes are regulated by the redox state of the cell or the organelle where the pathway occurs (Glaring *et al*., 2012, Lepisto *et al*., 2013). It is possible that GlgB does not behave like Arabidopsis BEs which are redox sensitive (Glaring *et al*., 2012), thus modifying the branching activity in the transgenic lines and explaining why it is not possible to restore a true WT phenotype.

Finally, it should be emphasized that the ability to form starch-like structures depends on the level of branching enzyme activity of GlgB. Indeed, β20, which possessed the highest GlgB activity, synthesizes polyglucans whose structure resembles more or less that of the residual insoluble glucans isolated from Arabidopsis DBE mutants (Wattebled *et al*., 2008). The β12 line, that had a GlgB activity intermediate between that of β1 and β20, displays an intermediate polyglucan-accumulating phenotype. Debranching enzyme activity level was estimated in the different lines by zymogram analysis (Figure S5). No obvious modification of isoamylase activity was detected in the GlgB-expressing plants compared to the wild type. Starch synthase activity was also evaluated with zymograms (Figure S6). SS1 and SS3 are the two major isoforms of starch synthases encompassing over 90% of the elongating activity measured in *Arabidopsis* leaf extracts (Szydlowski *et al*., 2011). Starch synthase activities were unmodified in the *be2 be3* mutant (Dumez *et al*., 2006). Both SS1 and SS3 activities were reduced in β1 compared to WT. The activity of SS1 was slightly higher in β12 and β20 compared to WT whereas SS3 was, to some extent, reduced. At this stage it is unclear why starch synthase activity is modified in the transgenic lines, especially in β1. Nevertheless, despite possessing the lowest starch synthase activity, β1 is the GlgB expressing line accumulating the highest amount of insoluble polyglucans. Thus the remaining SS activity in β1 appears in be in excess of that needed to allow the synthesis of the insoluble polyglucan, and does not seem to be a limiting factor.

Thus, in a context of non-limiting starch synthase activity, the accumulation of the insoluble and soluble polyglucans in β20 occurs under circumstances of over-branching which cannot be balanced by the debranching activity of endogenous isoamylases and pullulanase. This results in a higher branching degree (7.3%) and a very low crystallinity index (13%) of the insoluble polyglucans in β20. Such increase of the branching degree of the polyglucan produced in plants after the expression of *E. coli* GlgB has already been described in potato and rice (Kortstee *et al*., 1996, Kim *et al*., 2005). In both cases, the number of branch points of amylopectin was significantly increased. However, because endogenous BE activity was not knocked down in these plants the actual contribution of GlgB for the synthesis of the highly branched starch was impossible to determine.

Our result suggests that a finely tuned balance between branching and debranching activities is probably acting *in planta* to control α(1→6) linkage placement and number, and consequently to allow the formation of starch or starch-like polyglucan. Figure 10 is an attempt to model our interpretation of the results generated in this work. As a model, it is likely a simplified view of reality (for instance starch synthases are not included in the model) and was conceptualized from the results of the expression of a bacterial enzyme in plants. However, we suggest that it could be generalized to any situation where the balance between BE and DBE activity is compromised. For instance, in rice, the endogenous BEIIb isoform was expressed in a BEIIb-defective mutant by expression of the corresponding structural gene (Tanaka *et al*., 2004). Overexpression of the protein and of the corresponding activity was obtained in one of the transgenic lines leading to the accumulation of significantly higher amount of soluble polyglucan compared to the wild type (almost three times more). This was accompanied by enrichment in DP<15 glucans of amylopectin and alteration of the crystalline structure of starch.

**Figure 10:**
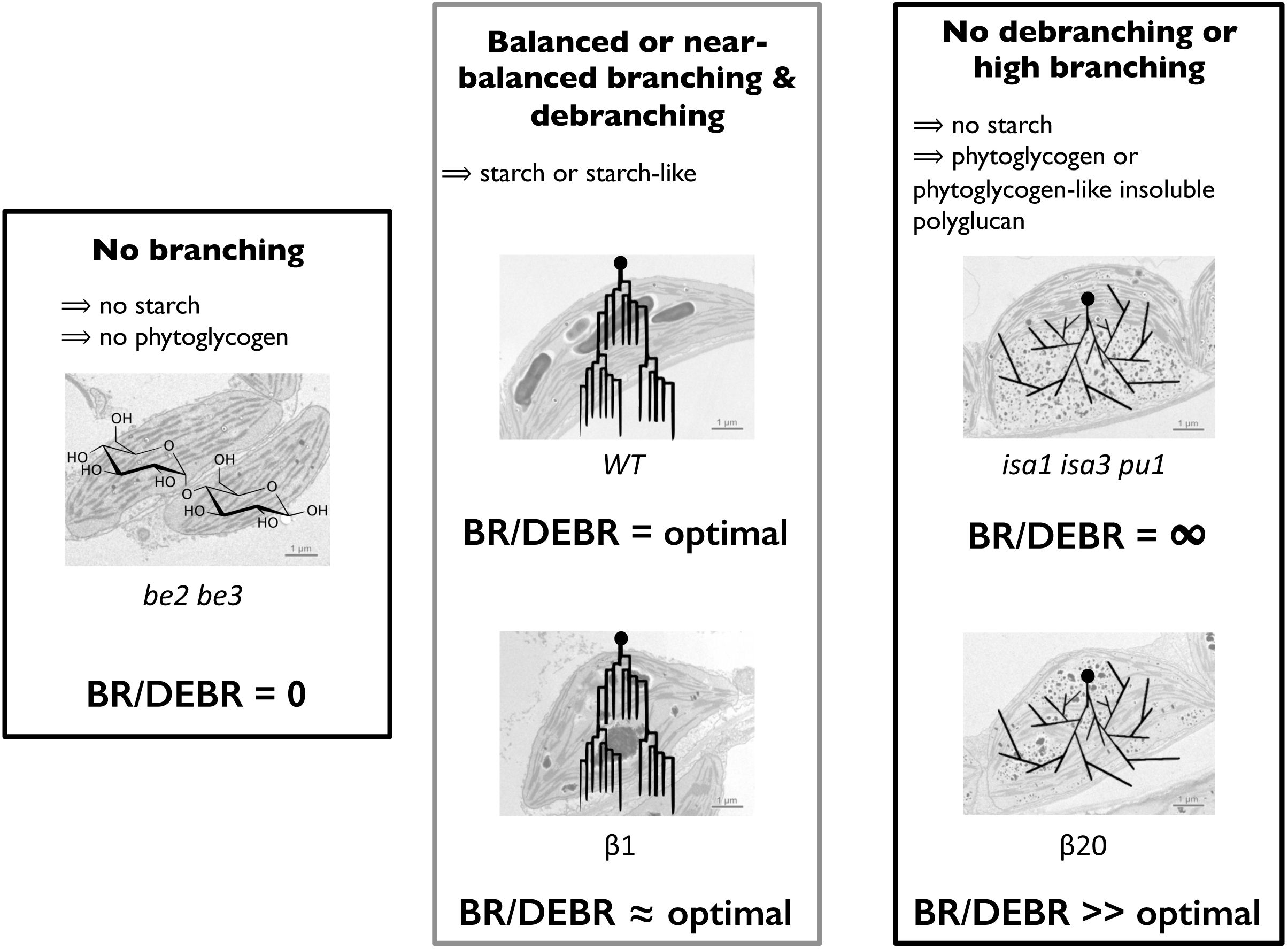
Starch or starch-like polyglucan synthesis occurs thanks to a balanced branching / debranching activity ratio assuming that starch synthase activity is not limiting whatever the branching/debranching ratio. Left panel: in the absence of branching activity (*be2 be3* double mutant), starch or phytoglycogen synthesis is impossible and is substituted by maltose accumulation; the BR/DEBR ratio is null. Middle panel: when the ratio BR/DEBR activity is balanced (WT) or near-balanced (β1), starch or starch-like polyglucan synthesis is promoted. In the case of β1, the slight structural modification of starch is due to the expression of the bacterial glycogen-branching enzyme that has different catalytic and regulatory properties. Right panel: in the absence of debranching activity (such as in *isa1 isa3 pu1;* BR/DEBR is infinite) or when branching activity is extremely high (such as in β20; BR/DEBR >> optimal) phytoglycogen or phytoglycogen-like insoluble polyglucan is synthesized respectively. Superimposed drawings correspond to (from left to right): maltose, amylopectin, phytoglycogen; black points depict the reducing end.

Nevertheless expression of GlgB did not restore the synthesis of true WT starch, possibly because GlgB activity cannot be regulated in the same way as endogenous BEs. Protein complexes and thus other enzyme activities (starch synthases for instance) might be affected by the lack of endogenous BEs, leading to the synthesis of structurally modified polyglucan as frequently observed in mutants defective for enzymes of the pathway.

## ACKNOWLEDGEMENTS

The authors thank Dr. Nicolas Szydlowski for fruitful discussions and critical reading of the manuscript, the NanoBio-ICMG Electron Microscopy Platform (Grenoble) and gratefully acknowledge the financial support of Agence Nationale de la Recherche (contract # ANR-11-BSV6-0003) and the Natural Sciences and Engineering Research Council of Canada (Team Discovery Grant, number 435781, MJE, IJT).

## SUPPORTING INFORMATION

Supporting Figure S1: **Description of the approach employed for the synthesis the GlgB chimeric sequence used for expression in *A. thaliana.***

Supporting Figure S2: **Comparison of mature plant size.**

Supporting Figure S3: **Iodine-staining phenotype of individual plants harvested at different time points.**

Supporting Figure S4: **Chain length distribution profiles of water-soluble polyglucans.**

Supporting Figure S5: **Zymogram analysis of starch and p-limit dextrins hydrolyzing enzymes from transformed plants expressing GlgB.**

Supporting Figure S6: **Zymogram analysis of soluble starch synthases activities.**

Supporting Figure S7: **Structure of polysaccharides produced in β5 and β6 transformant lines.**

Supporting Figure S8: **Leaf glucan contents and granule morphology.**

**Supplemental Figure S1:**
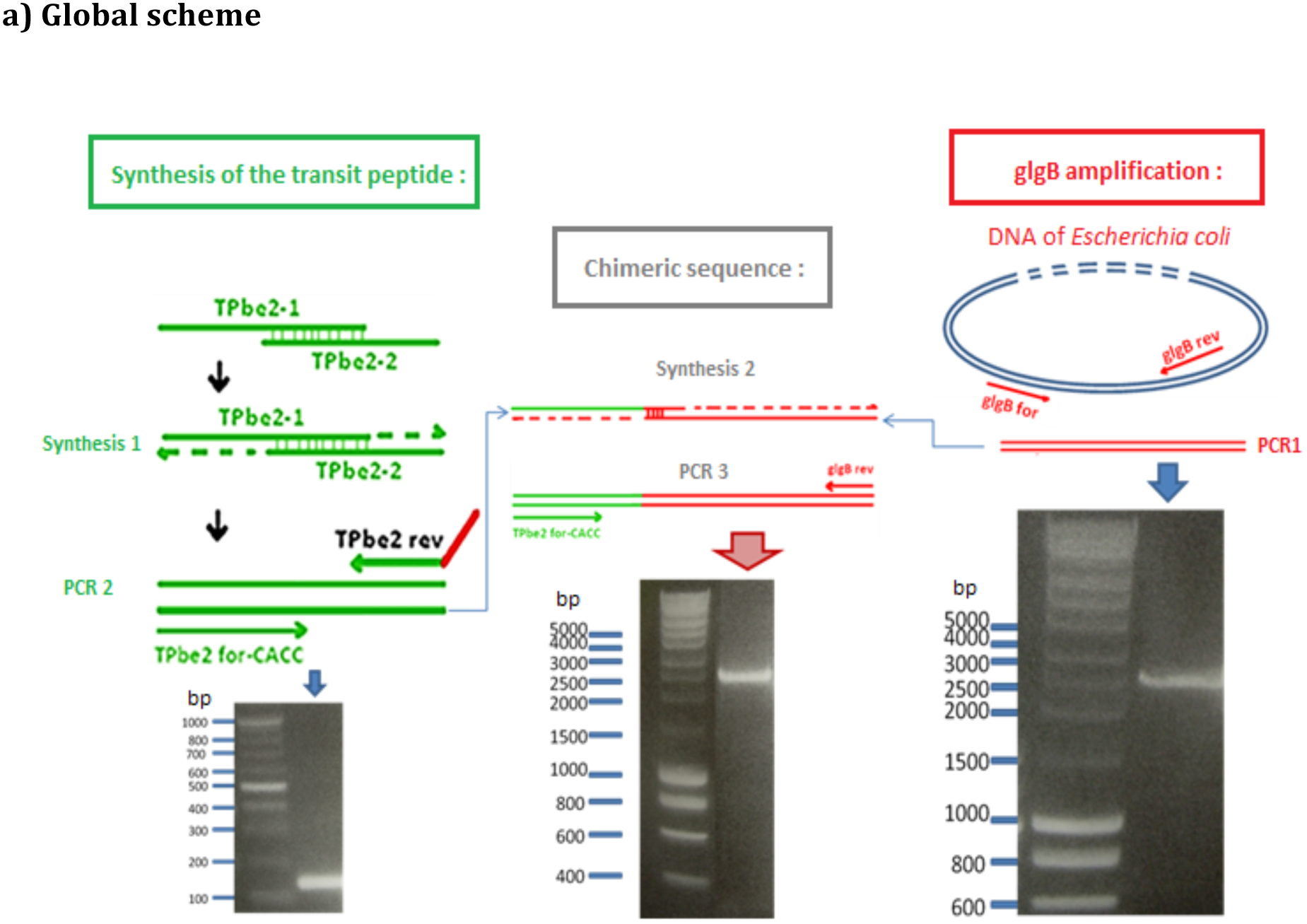
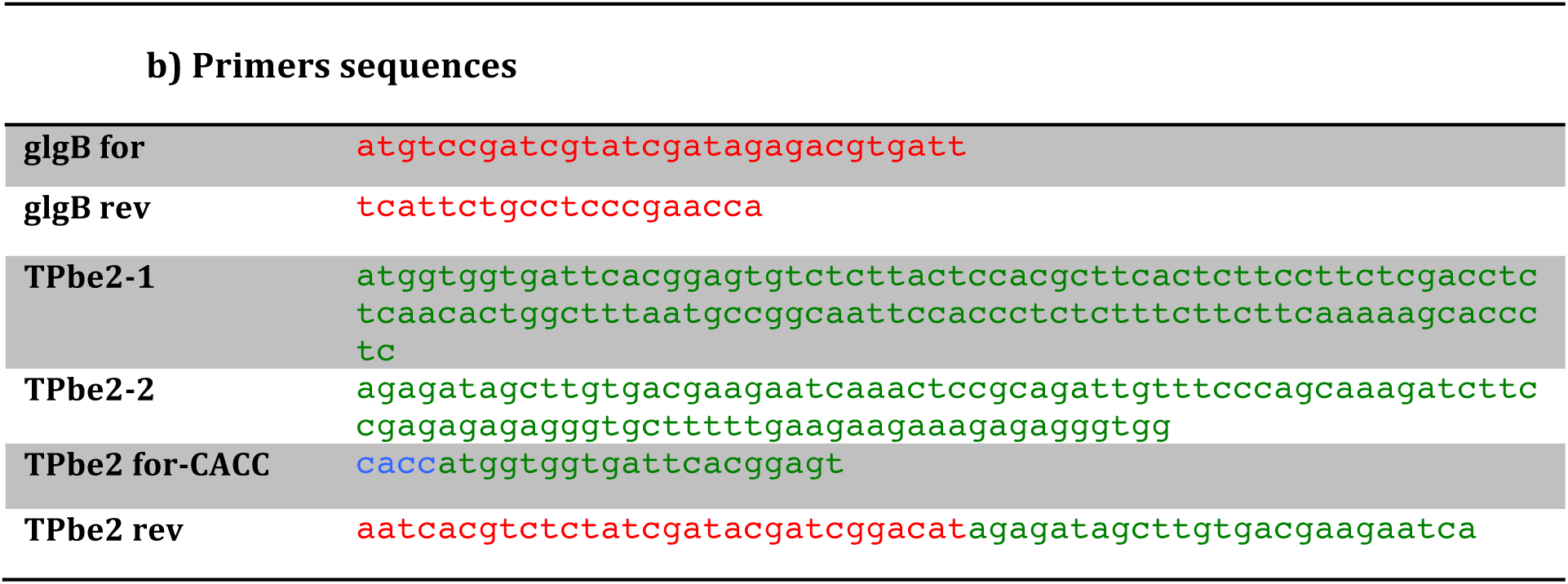
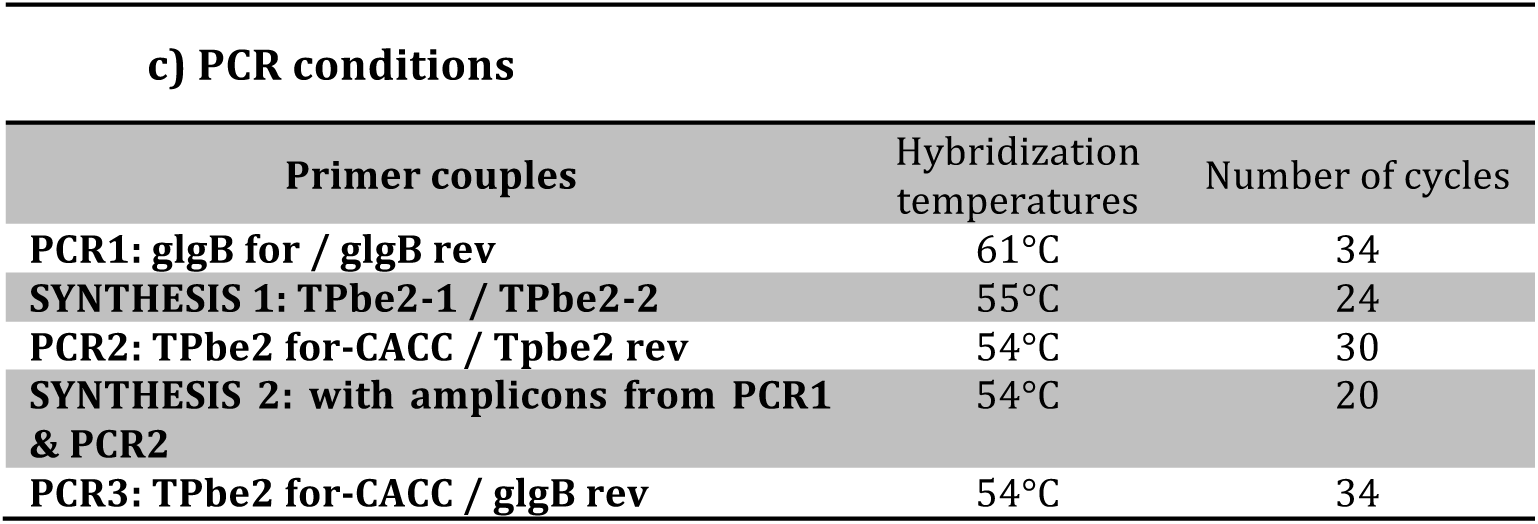
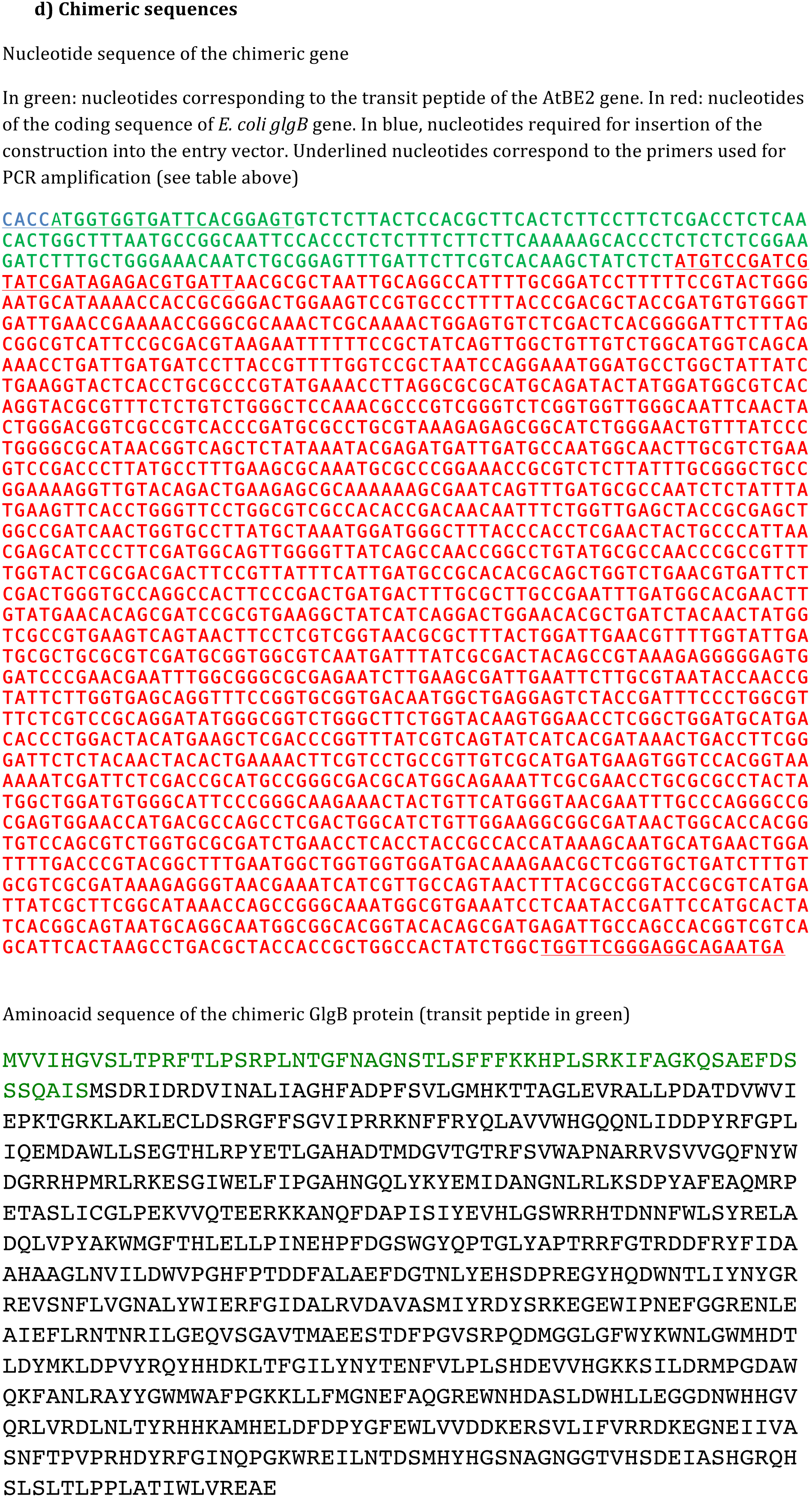
Description of the approach employed for the synthesis the GlgB chimeric sequence used for expression in *A. thaliana*. The chimeric sequence used to transform the *be2 be3* double mutant is composed of the nucleotide sequence of the transit peptide of the plant gene AtBE2 (At5g03650) and the E. coli *glgB* nucleotide sequence (M13751). a) Global scheme of the method is presented. b) Primers nucleotide sequences. Nucleotides in green correspond to the AtBE2 transit peptide sequence and nucleotides in red correspond to the coding sequence of *glgB.* Nucleotides “CACC” highlighted in blue are required for insertion of the sequence in the pENTR/D-TOPO vector. (c) Hybridization temperatures used for each couple of primers and number of cycles used during amplification. (d) Nucleotide and amino acid sequences of the chimeric gene.

**Supplemental Figure S2:**
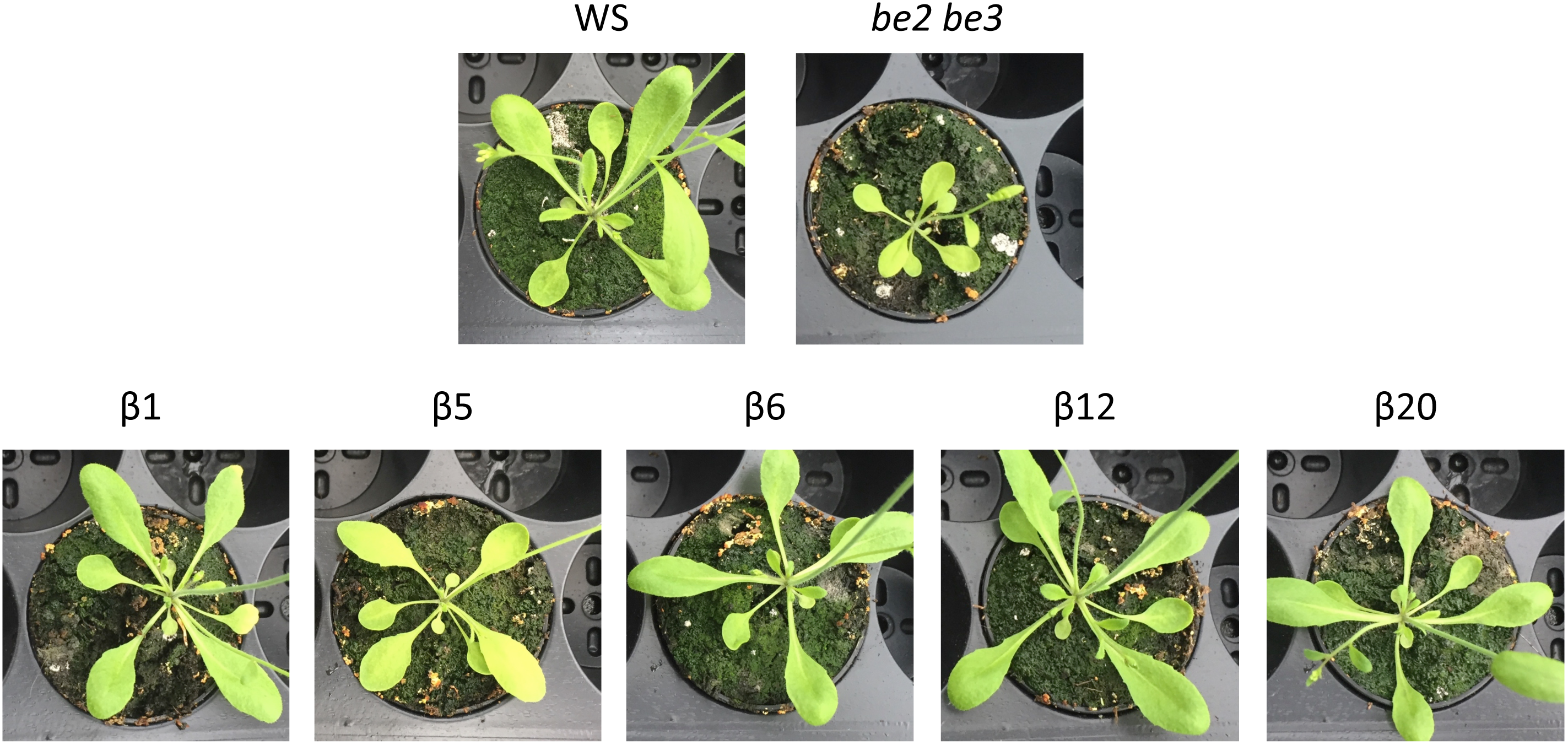
comparison of mature plant size. Seeds were sown on peat-based compost and plants were grown in greenhouse for 16h in the light at 21°C and 8h in the dark and 16°C. Pictures were taken 3 weeks after germination. The *be2 be3* double mutant has a strong growth retardation phenotype and plants are much smaller than wild type (Dumez et al., 2006) whereas all transformed plants expressing *E. coli* GlgB branching enzyme have a phenotype close to that of the wild type reference.

**Supplemental Figure S3:**
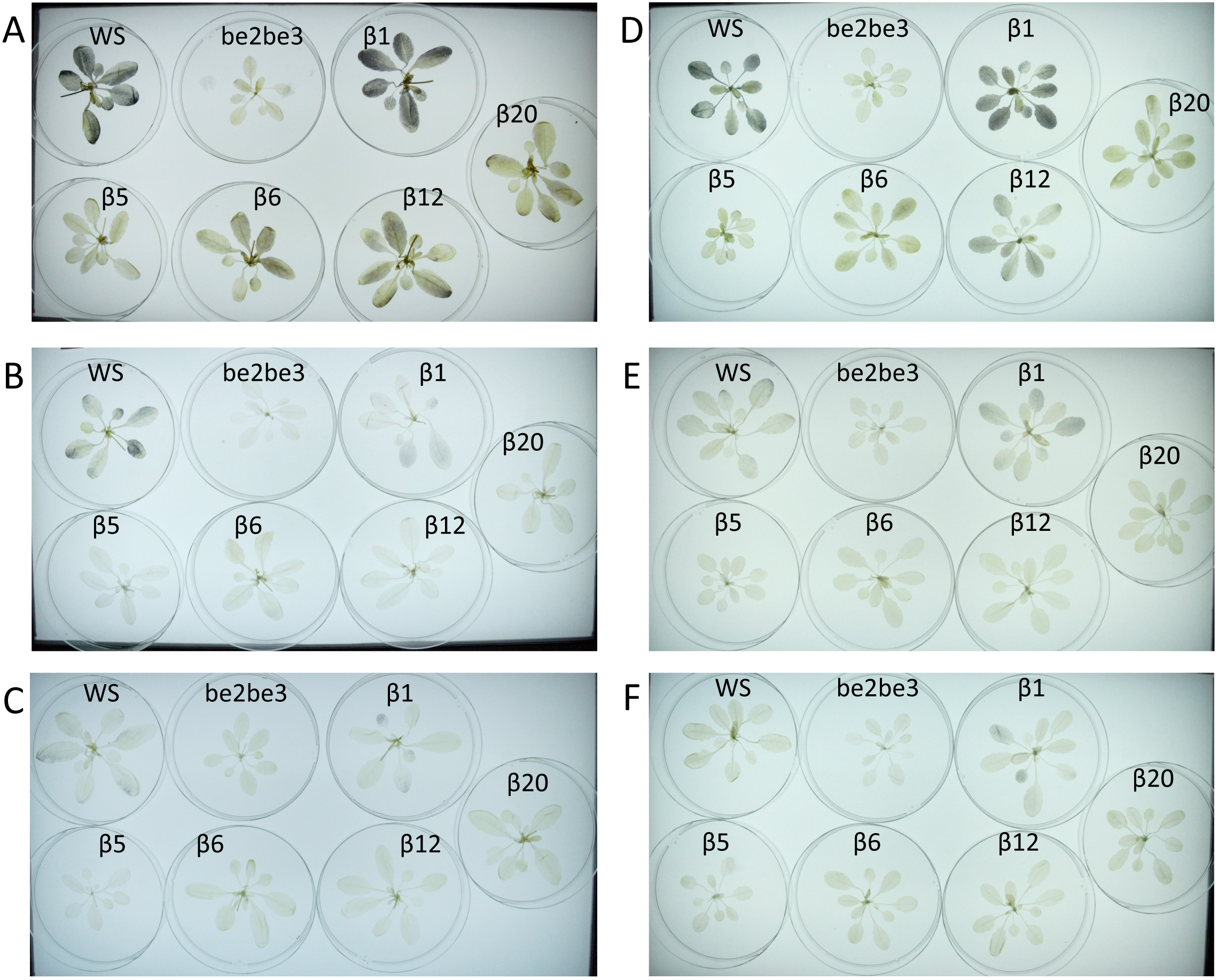
Iodine-staining phénotype of individual plants harvested at different time points. Plants were cultivated in growth cabinet in 16-h light; 8-h dark (A, B, C) or in 12-h light; 12-h dark (D, E, F) regime at 23°C during the day and 20°C during the night and 100 μE.m^−2^.s^−1^ irradiance. Plants were harvested at the end of light period (A and D) or at the end of the dark period (B and E). In panels C and F plants were left 24 more hours in the dark. All plants were destained in hot ethanol and stained with iodine solution. At the end of the dark period (Panels B and E) polysaccharide degradation is either complete (β5, β6, β12, β20;) or similar to the wild type line (β1). After an extend of 24 hours in darkness, all lines appear colorless.

**Supplemental Figure S4:**
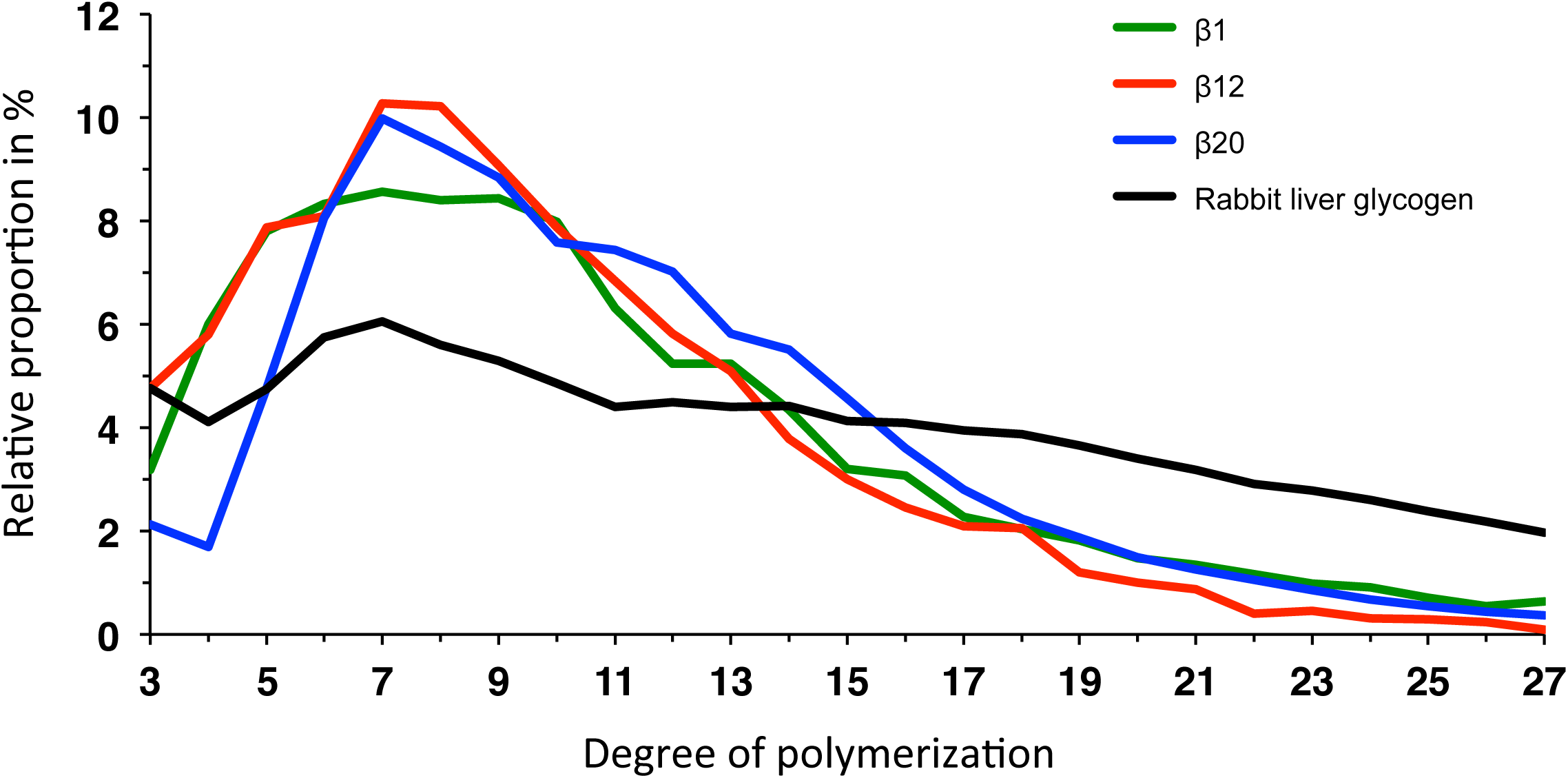
Chain length distribution profiles of water soluble polyglucans. purified from GlgB-expressing lines β1 (green), β12 (red) and β20 (blue) compared to that of the rabbit liver glycogen (black). After purification, water soluble polyglucans were debranched by a mix of bacterial isoamylase and pullulanase. The branched products were analyzed by HPAEC-PAD. The relative proportion of each glucan is plotted versus their degree of polymerization. Values are the mean of two analysis carried out with soluble polyglucans extracted of plants cultivated independently.

**Supplemental Figure S5:**
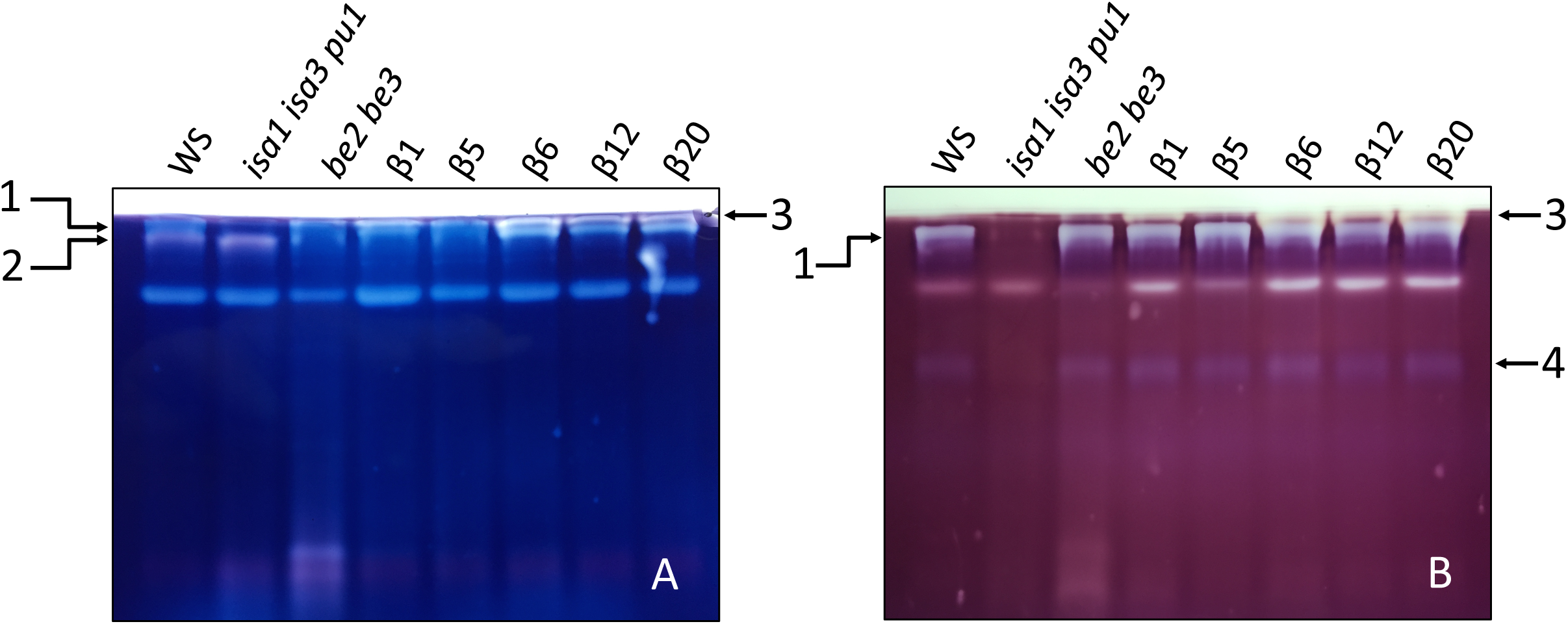
Zymogram analysis of transformed plant expressing GlgB. Three to four leaves of 3 week-old plants were harvested at mid-day for soluble extract preparation. The equivalent of 50 μg of proteins were loaded on polyacrylamide gel impregnated with either 0.3% of potato soluble starch (A) or 0.3% of maize β-limit dextrins (B). After migration (2 h at 4°C, 15 mA / gel), the gels were incubated overnight at room temperature in a buffer composed of: Tris 25 mM; Glycine 192 mM; MgCl_2_ 1 mM; CaCl_2_ 1 mM; DTT 10 mM. After incubation, enzymes that modified the substrate were revealed by soaking the gels in iodine solution (1% KI [w/v]; 0.1% I_2_ [w/v]). The gels were rinsed in water before taking pictures. WS is the wild type reference (*Wassilewskija*) and all other plants are in the same genetic background. *be2 be3* corresponds to a double mutant lacking endogenous branching enzymes. β1, β5, β6, β12, and β20 are *be2 be3* mutants expressing different levels of *E. coli* GlgB. Blue Band labeled 1 and red band labeled 2 were previously determined as Iso1 and BE2 respectively (Dumez et al., 2006; Wattebled et al., 2008). The pale pink band labeled 3 with an extremely low mobility, present only in transformed plants, corresponds to GlgB. Band 4 is the pullulanase (Wattebled et al., 2008).

**Supplemental Figure S6:**
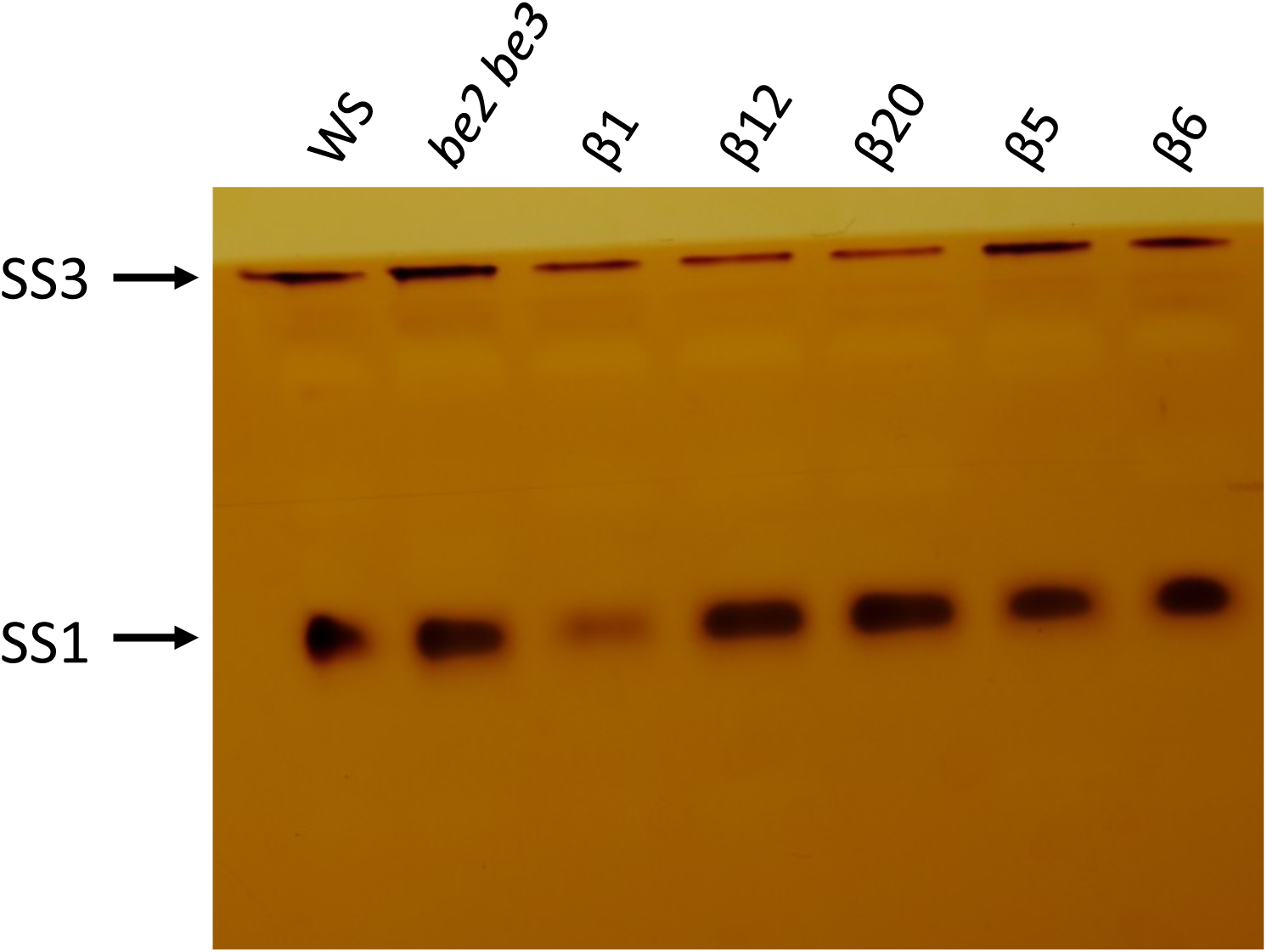
Zymogram analysis of soluble starch synthases activities. An equal amount of proteins from leaf crude extracts was loaded on native PAGE containing 0.3% (w/v) of rabbit liver glycogen. Approximately 2h30 of migration in native conditions (Tris-glycine buffer 1X, 4°C) was applied. After incubation in an appropriate buffer for starch synthase activity (Glycyl-glycine 66 mM (pH 7.5); (NH_4_)_2_SO_4_ 66 mM; MgCl_2_ 1 mM; β-mercaptoethanol 3.3 mM; ADP-glucose 1.2 mM), the gel was washed 6 times into water to remove β-mercaptoethanol and further incubated in iodine solution until revelation of bands of activity. This zymogram is representative of three biological independent experiments.

**Supplemental Figure S7:**
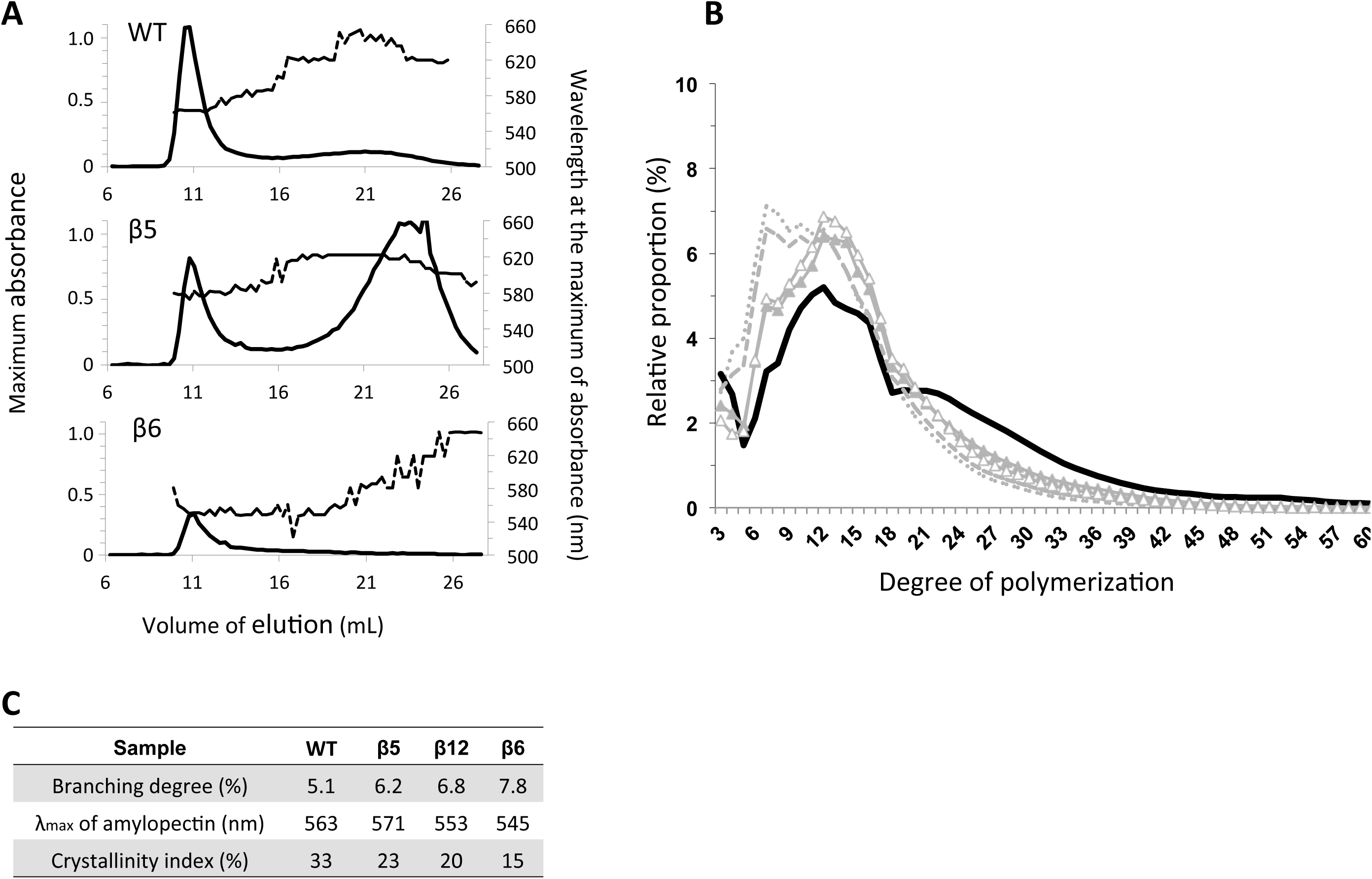
Structure of polysaccharides produced in β5 and β6 transformant lines. **(A) Fractionation of insoluble polyglucans by size exclusion chromatography.** Dashed lines indicate the wavelength (nm) of the iodine-polyglucan complex at the maximum of absorbance (right Y-axis). **(B) Chain length distribution of insoluble polyglucans.** WT: continuous black line; β1: closed grey triangles; β5: open grey triangles; β6: dotted grey line; β20: discontinuous grey line. **(C) Structural parameters of polysaccharides produced in β5 and β6 tranformant lines.** The branching degree was calculated according to Szydlowski et al. (2011).

**Supplemental Figure S8:**
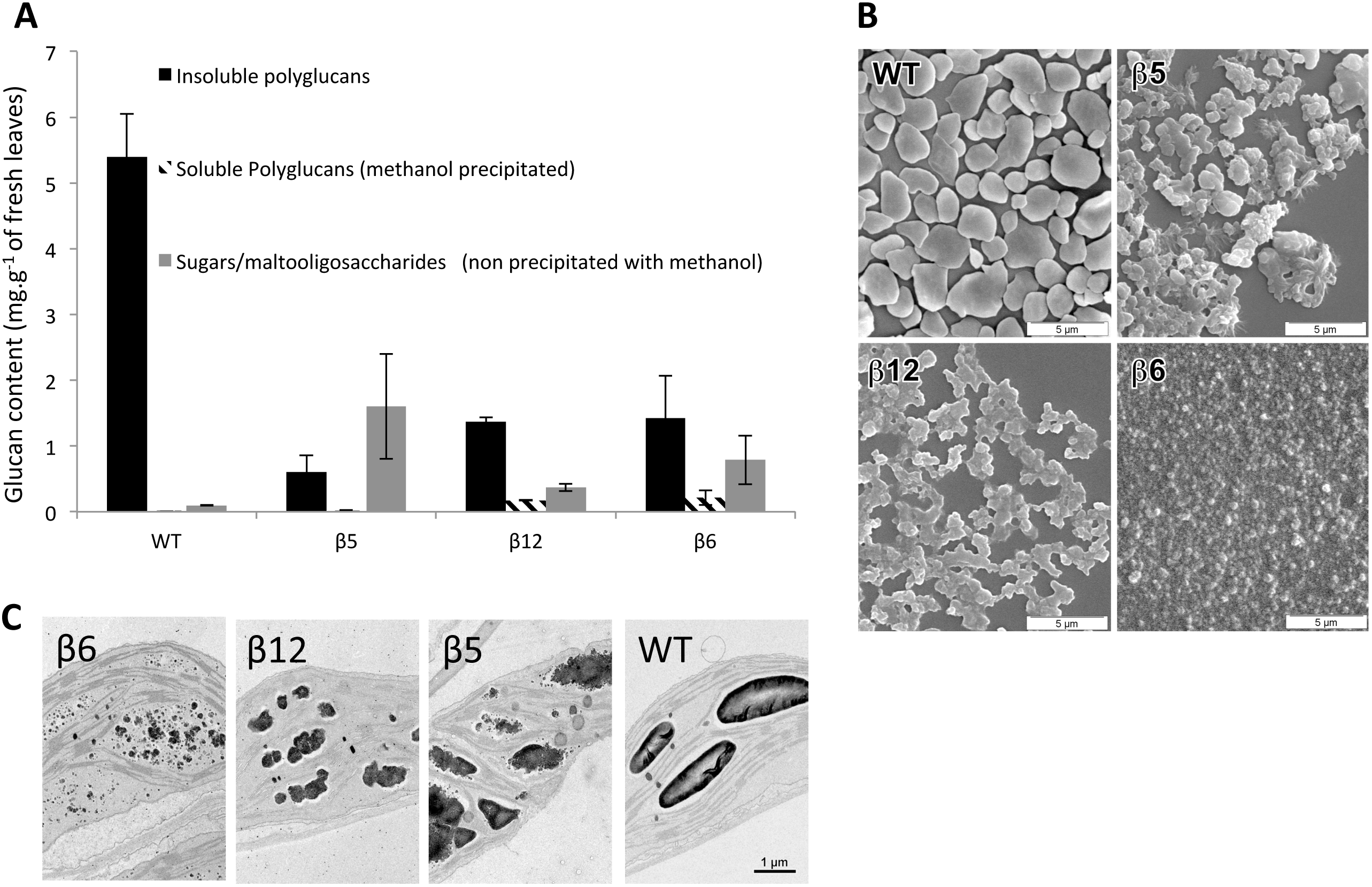
Leaf glucan contents and granules morphology. A: For glucan content the average of three independent cultures are presented. Vertical thin bars are the standard deviation of three independent biological replicates. B: Scanning electron microscopy images of purified insoluble polyglucans. C: Transmission electron microscocopy images of leaf chloroplasts positively stained with PATAg.

